# Ubiquitin-dependent degradation of p21^Waf1/Cip1^ is mediated by AMBRA1 to limit DNA replication stress

**DOI:** 10.1101/2025.07.28.666126

**Authors:** Giacomo Milletti, Giulia Cadeddu, Alba Adelantado-Rubio, Caterina Ferraina, Kristina Keuper, Sebastian Munk, Cristiano De Stefanis, Denise Quacquarini, Sabrina Rossi, Angela Mastronuzzi, Franco Locatelli, Paolo Grumati, Valentina Cianfanelli, Francesca Nazio, Jiri Bartek, Apolinar Maya-Mendoza, Francesco Cecconi

**Affiliations:** DNA Replication and Cancer Group, Danish Cancer Institute, Copenhagen, Denmark; Hematology/Oncology, Cell Therapy, Gene Therapies and Hemopoietic Transplant, Bambino Gesù Children’s Hospital, IRCCS, Rome, Italy; Department of Biology, University of Rome Tor Vergata, Rome, Italy; Department of Woman and Child Health and Public Health, Gynecologic Oncology Unit, Fondazione Policlinico Universitario A. Gemelli IRCCS, Rome, Italy; Research Facilities, Bambino Gesù Children’s Hospital IRCSS, Rome Italy; Pathology Unit, Bambino Gesù Children’s Hospital IRCSS, Rome Italy; Department of Health Science and Public Health, Catholic University of the Sacred Heart, Rome, Italy; Telethon Institute of Genetics and Medicine, Pozzuoli, Naples, Italy; Department of Science, University “ROMA TRE”, 00146, Rome, Italy; Immunology Research Area, Innate Lymphoid Cells Unit, Bambino Gesù Children’s Hospital (IRCCS), Rome, Italy; Genome Integrity Group, Danish Cancer Institute, Copenhagen, Denmark; Science for Life Laboratory, Division of Genome Biology, Department of Medical Biochemistry and Biophysics, Karolinska Institutet, S-171 21 Stockholm, Sweden; Cell Stress and Survival Group, Center for Autophagy, Recycling and Disease (CARD), Danish Cancer Institute, Copenhagen, Denmark; Università Cattolica del Sacro Cuore and Fondazione Policlinico Universitario Agostino Gemelli IRCCS, Rome, Italy

## Abstract

The cell cycle is a fundamental process that orchestrates the events that lead to cell replication and division. AMBRA1 is a scaffold factor that interacts with the E3 ubiquitin ligase CRL4-DDB1 complex to regulate the stability of cyclin D-type proteins, key regulators of the G1/S phase transition. However, how cells complete S-phase and coordinate the speed of DNA synthesis is still to be fully elucidated. Here we demonstrate that AMBRA1 affects the turnover of p21^Waf1/Cip1^ and p27^Kip1^, by coupling the CRL4-DDB1 complex directly to these proteins. In the absence of AMBRA1, the increased stability of p21^Waf1/Cip1^, rather than p27^Kip1^, resulted in the accumulation of replication stress by leaving under-replicated DNA during the S phase. We also found that AMBRA1-depleted cells with consequent high p21^Waf1/Cip1^ are sensitive to the inhibition of lagging-strand DNA synthesis. Of clinical relevance, low levels of AMBRA1 in Sonic-Hedgehog medulloblastoma correlate with a worse prognosis, suggesting that AMBRA1 expression levels could be an informative criterion for patient stratification and treatment.

## Introduction

Cells must coordinate the speed of DNA synthesis with the length of the cell cycle to ensure genome integrity^1^. The length of the cell cycle is determined by growth factors and metabolic signals that are ultimately integrated by cyclins (including cyclins A, B, C, D and E) and cyclin-dependent kinases (CDKs). Specific cyclins modulate transitions through the cell cycle by synthesis and degradation processes. D-type cyclins in complex with CDK4/6 drive the transition from G1 to S phase. The accumulation of cyclin D1 due to aberrant regulation results in a fast cell cycle, fast speed of DNA replication fork progression and genome instability^2^. Several other proteins and post-translational modifications act in concert to regulate each phase of the cell cycle, classical examples of those are p21^Waf1/Cip1^ and p27^Kip1^ (hereafter referred as p21 and p27) ^3–6^. p27 and p21 inhibit the activity of CDK2-containing complexes and serve as assembly factors for cyclin D–CDK4/6^7,8^. The latter role was demonstrated by the observation that mouse fibroblasts lacking p27 and p21 fail to assemble cyclin D–CDK4/6 complexes^9^. The Activating Molecule in Beclin 1-Regulated Autophagy (AMBRA1) acts as a substrate receptor for CRL4-mediated degradation of D-type cyclins, restraining G1/S transition and therefore preserving genome integrity^2,10^. Cullin-RING finger ligases (CRLs) are the largest family of ubiquitin ligases in eukaryotic cells, regulating diverse cellular processes. Unlike other cullins, CUL4 (CUL4A and CUL4B) employs DDB1, a WD40-like repeat-containing adaptor, to assemble the CRL4-DDB1 complex^11^. This E3 ubiquitin ligase governs protein turnover through ubiquitination, playing a crucial role in DNA replication^12,13^, repair, and cell cycle control, thereby ensuring cellular homeostasis. Similarly to cyclins, numerous E3 ubiquitin ligases have been implicated in the recognition of p27 and p21; however, the full regulatory network governing their turnover remains to be fully elucidated. Understanding these mechanisms is essential for deciphering cell cycle control and its impact on genomic stability.

Growing evidence highlights the crucial role of AMBRA1 in cancer development. Notably, AMBRA1 expression varies among different medulloblastoma (MB) subgroups^14^, with the lowest levels observed in Sonic Hedgehog-MB (MB-SHH), a subtype characterized by high genomic instability^15^. This instability is associated with a replication stress (RS) phenotype similar to that caused by AMBRA1 depletion^16^. Moreover, a Sleeping Beauty (SB) transposon mutagenesis analysis identified AMBRA1 as a novel genetic driver of embryonal brain tumors primed by p53 loss^17^, a condition previously observed also in lung cancer^2^.

Here, we show that AMBRA1 coordinates the degradation of p27 and p21, besides that of D-type cyclins, thus impacting the balance and activity of CDK4/6 and CDK2 complexes. Therefore, AMBRA1 modulates genomic DNA synthesis indirectly by maintaining adequate levels of p21. The accumulation of p21 results in RS due to the acceleration of fork progression leaving under-replicated DNA. Mechanistically, the excess of p21 binds to PCNA interfering with the binding of other PCNA-interacting factors. Furthermore, we elucidate the role of AMBRA1 as a tumor suppressor in p53-null MB-SHH cell lines, where the permissive environment generated by p53 mutation enables AMBRA1-depleted cells to cope with elevated RS.

Last, our data shed light on the clinically relevant phenomenon of MB-SHH resistance to CDK4/6 inhibitors chemotherapy^18,19^, but sensitivity to lagging DNA synthesis inhibition, thus heralding potential innovative therapeutic avenues for this lethal disease.

## Results

### AMBRA1 regulates the stability of p27 and p21 in MB-SHH cell lines

To gain deeper insight into the impact of AMBRA1 loss on cellular physiology, we analyzed data from the Cancer Dependency Map Portal^20^ to identify genes exhibiting co-dependencies with AMBRA1 knockout (AMBRA1-KO). Gene Effect correlations revealed a potential involvement of AMBRA1 in p21 and p27 degradation (Fig. 1a). Given its potential significance, we examined p21 and p27 protein levels in AMBRA1-KO cells, using cyclin D1 and D3 as controls, both before and after AMBRA1 reconstitution. Notably, AMBRA1 depletion led to the accumulation of p21 and p27, a phenotype that was reversed by AMBRA1 overexpression (Fig. 1b). We have previously shown that AMBRA1 is differentially expressed across the molecular subtypes of MB with the highest expression in Group 3 and WNT and the lowest in SHH^14^, making MB an ideal model to study the effect of different expression levels of AMBRA1 in cell pathophysiology. Analysis of available MB-SHH datasets (deposited at http://r2.amc.nl) revealed that AMBRA1 low expression correlates with reduced survival and that its mRNA levels are downregulated in this tumor context compared to normal cerebellar cells (Fig. 1c, d). Using a patient cohort of 14 MB-SHH cases, immunohistochemical analysis for the levels of AMBRA1, cyclin D1 and p21, found that these proteins inversely correlate, suggesting that AMBRA1 may be involved in the degradation of the cyclin D1 and p21; in fact, in tumors with a high expression of AMBRA1, cyclin D1 and p21 were downregulated (Fig. 1e and Extended Data Fig. 1a).

**Figure 1.**
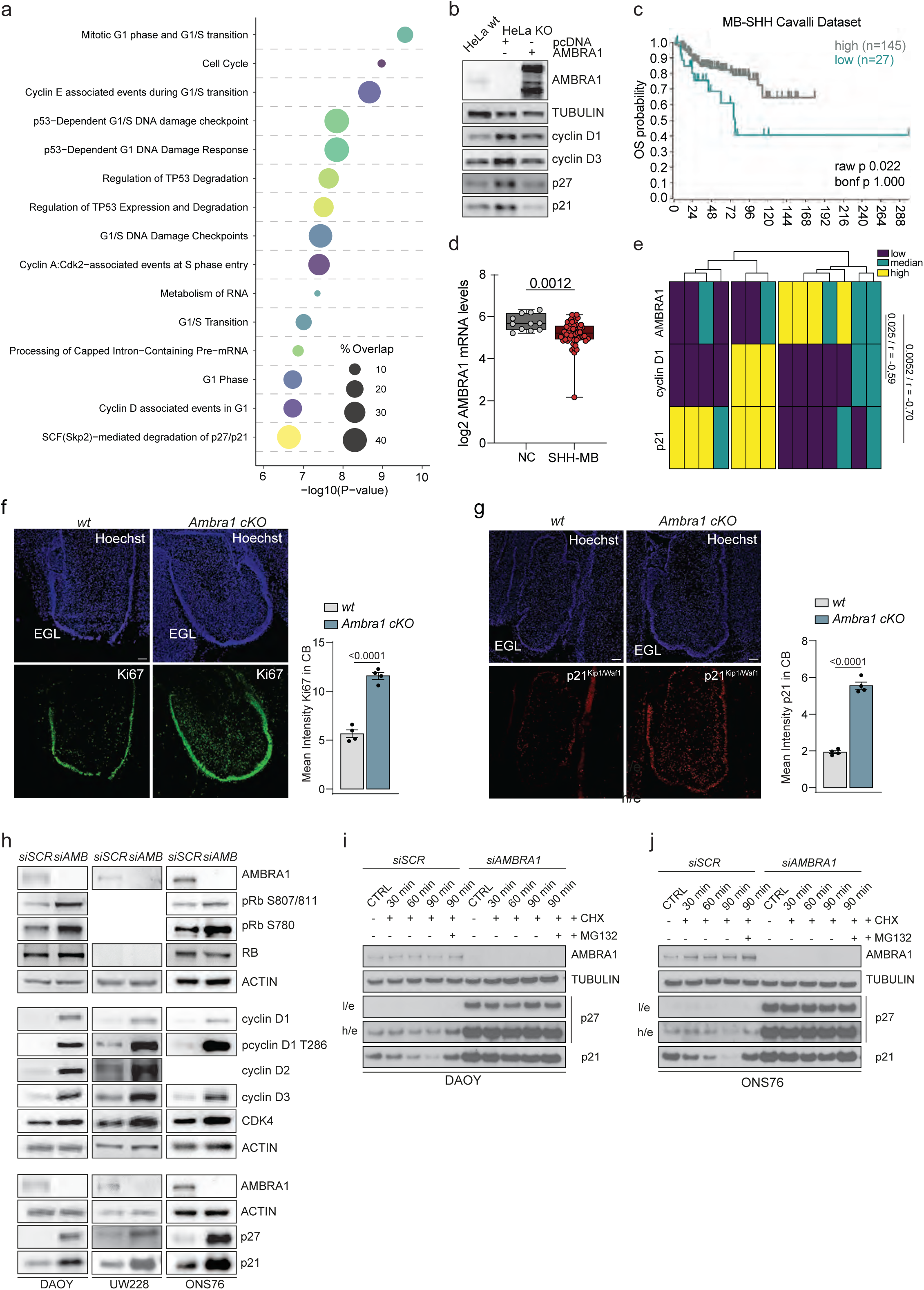
AMBRA1 regulates p21 and p27 stability in MB-SHH cell lines. **a**, Bubble plot of pathway enrichment analysis for AMBRA1 Top500 co-dependencies from DepMap CRISPR KO database. **b**, Immunoblot (IB) of the indicated proteins in wt or *AMBRA1*-KO HeLa cells, reconstituted or not with a plasmid overexpressing AMBRA1 (n=3). **c**, Kaplan– Meier analysis for MB SHH patients according to AMBRA1 expression levels in Cavalli dataset. Low AMBRA1 n = 27; High AMBRA1 n = 145. Expression cut-off 239.500. **d**, Boxplot for mRNA log2 expression of AMBRA1 derived from the publicly available dataset Pomeroy (204 patients, MAS 5.0 normalized, u133a platform), presented as mean value ±SD. NC, normal cerebellum = 11; MB-SHH = 54. **e**, Heatmap of immunohistochemistry (IHC) staining levels for the indicated proteins in our cohort of MB-SHH patients. The intensities were ranked in three grades, shown as negative or weak positive (*low*), median positive (*median*), and strong positive (*high*). **f-g,** Sagittal sections from wild-type and Ambra1 cKO E18.5 embryos stained for Ki67 (**f**) or p21(**g**) and Hoechst (n=4), with their respective intensity quantification of the cerebellar area. EGL, external granule layer. Scale bars 100 µm. **h**, IB for the indicated proteins in three MB-SHH cell lines: DAOY, UW228 and ONS76 in control or AMBRA1-depleted condition with small interfering RNA (siRNA). Rb and cyclin D2 were not detected due to their absence of expression in the specific cell lines used. **i**-**j**, IB of the indicated proteins in DAOY and ONS76 cells depleted for AMBRA1 with siRNA and treated with 100 µg/ml cycloheximide (CHX) and/or 10 µM MG132 for the indicated time points (n=3). Unless otherwise stated data are presented as mean value ±SEM and n refers to biological independent samples. Data were analyzed using log-rank test (**c**), Welch’s two-tailed unpaired t-test (**d**), Pearson correlation (**e**) and Mann-Whitney t-test (**f**-**g**).

The intricate shaping of the cerebellum relies on a meticulous regulation of the cell cycle, directing the fate decisions of individual cells during differentiation and stem cell functions^21^. MB, as an embryonal pediatric cancer, stems from aberrant cerebellar development. To investigate the impact of *Ambra1* absence on cerebellar development *in vivo*, an *Ambra1^flox/flox^*/*Nestin-Cre* mouse model was used. Cre recombinase activation led to *Ambra1* deletion, resulting in a marked increase in cerebellar anlage proliferation at E13.5 (Extended Data Fig. 1b) and correlating with elevated expression of D-type cyclins (Extended Data Fig. 1c). Later in development (E18,5), the newly formed cerebellum displayed a more proliferative external granule layer (EGL) (Fig. 1f), a region populated with granule cell precursors (GCPs). Indeed, *Ambra1* depletion resulted in high levels of cyclin D1, p21, and phospho-Rb (S807-811) in this region (Fig. 1g and Extended Data Fig. 1d). Altogether, these results underscore the critical role of AMBRA1 in normal cerebellar development and suggest its potential significance in MB.

To elucidate the molecular processes regulated by AMBRA1 in MB-SHH, we used three available cell lines isolated from this tumor subtype, DAOY, UW228 and ONS76^22^. In AMBRA1-depleted condition, we observed an increase in phospho-Rb (S807-811) in DAOY and ONS76 cell lines, however, in the UW228 cell line, there were no detectable levels of either total or phosphorylated Rb. In all three cell lines, the reduction of AMBRA1 resulted in the accumulation of cyclins D1-3, together with higher CDK4, p21 and p27 (Fig. 1h). We reasoned that similarly to the augmented stability of cyclins D, AMBRA1 could modulate the degradation of p21 and p27. Indeed, upon AMBRA1 depletion, the inhibition of de novo protein synthesis using cycloheximide (CHX), coupled with protein degradation impairment by proteasome inhibition (MG132), showed a significant increase in the protein stability of both proteins in DAOY and ONS76 cells (Fig. 1i-j and Extended Data Fig 1e, f). Additionally, we fitted an exponential decay model to estimate the precise half-lives of p21 and p27 in both cell lines. Upon AMBRA1 silencing, we observed that p21 stability increased significantly - doubling in ONS76 cells (107’ vs 182’) and tripling in DAOY cells (81’ vs 251’). Alongside, the half-life of p27 in AMBRA1-depleted cells could not be determined within the selected time frame (controls - ONS76: 137’; DAOY: 56’), indicating a marked stabilization of both proteins (Extended Data Fig 1g). To extend these observations, p27 and p21 transcriptional levels were analyzed following AMBRA1 suppression (Extended Data Fig. 1h), revealing no consistent statistically significant pattern across the three cell lines. Taken together, these results show that AMBRA1 is a key regulator of cyclin D1, p27 and p21 stability in MB-SHH.

### AMBRA1-mediated degradation of p21 and p27 is both dependent and independent of cyclin D1

Overexpression of cyclin D1 has been reported to promote stabilization of p21 in a proteasome-dependent manner by competing for its proteasomal binding, rather than by reducing its ubiquitylation^23^. Consistent with this mechanism, we also observe that overexpressing the three cyclin D family members stabilizes p21 and p27 in our MB model cell lines (Extended Data Fig. 2a). Since the absence of AMBRA1 also resulted in cyclin D1 accumulation^2,10,24^, we hypothesized that AMBRA1 could affect p21 stability in a cyclin D-dependent manner. To this end, we generated cells with doxycycline-inducible cyclin D1 expression in both AMBRA1-wt and -KO cells and treated them with low UV irradiation, a well-known trigger of p21 degradation as part of the DNA damage response^3^. Immunoblot analysis confirmed that while cyclin D1 overexpression alone reduced the degradation of p21, its effect was less pronounced than that of AMBRA1-KO. Moreover, the combined treatment showed no additional impact beyond AMBRA1-KO alone (Fig. 2a). This implies the existence of an alternative mechanism independent of cyclin D1 for the degradation of p21 and presumably also for p27. To test this possibility, we used a recently developed PROTAC degrader for cyclin D1, MS28^25^. First, AMBRA1 protein degradation was induced using a Flag-tagged minimal auxin-inducible degron (2×Flag–mAID), followed by degradation of cyclin D1. The results showed that the cyclin D1 depletion did not affect the enhanced stability of p27 and p21 in the absence of AMBRA1 (Fig. 2b). To exclude that this phenotype was cell-cycle dependent, we engineered DAOY cells to constitutively express cyclin D1 (T286A), a mutant insensitive to ubiquitin-mediated degradation that is not able to interact with AMBRA1^2,10^. In this transgenic cell line, we collectively knockdown each of the endogenous D-type cyclins alone or in combination with AMBRA1 depletion. The experiments revealed that p21 and p27 were upregulated even in the absence of cyclin D, albeit to a lesser extent (Fig. 2c). Notably, silencing of cyclins D significantly attenuated the stabilization of p21 and p27 caused by AMBRA1 depletion, supporting a parallel cyclin-dependent regulatory mechanism. Furthermore, the overexpression of cyclin D1 did not impact the levels of ubiquitination of p27 or p21 (Extended Data Fig. 2b). Intriguingly, the accumulation of cyclin D1 due to AMBRA1 depletion was prevented in both MB cell lines upon p21 co-depletion, whereas p27 silencing had a minor and inconsistent effect across the two cell lines (Extended Data Fig. 2c). Moreover, the fluctuation of the protein levels in the tested experimental conditions was independent of the mRNA levels (Extended Data Fig. 2d). Overall, the results show a direct involvement of AMBRA1 in the regulation of p27 and p21 stability, a phenomenon that occurs at least in part independently of cyclin D1 levels.

**Figure 2.**
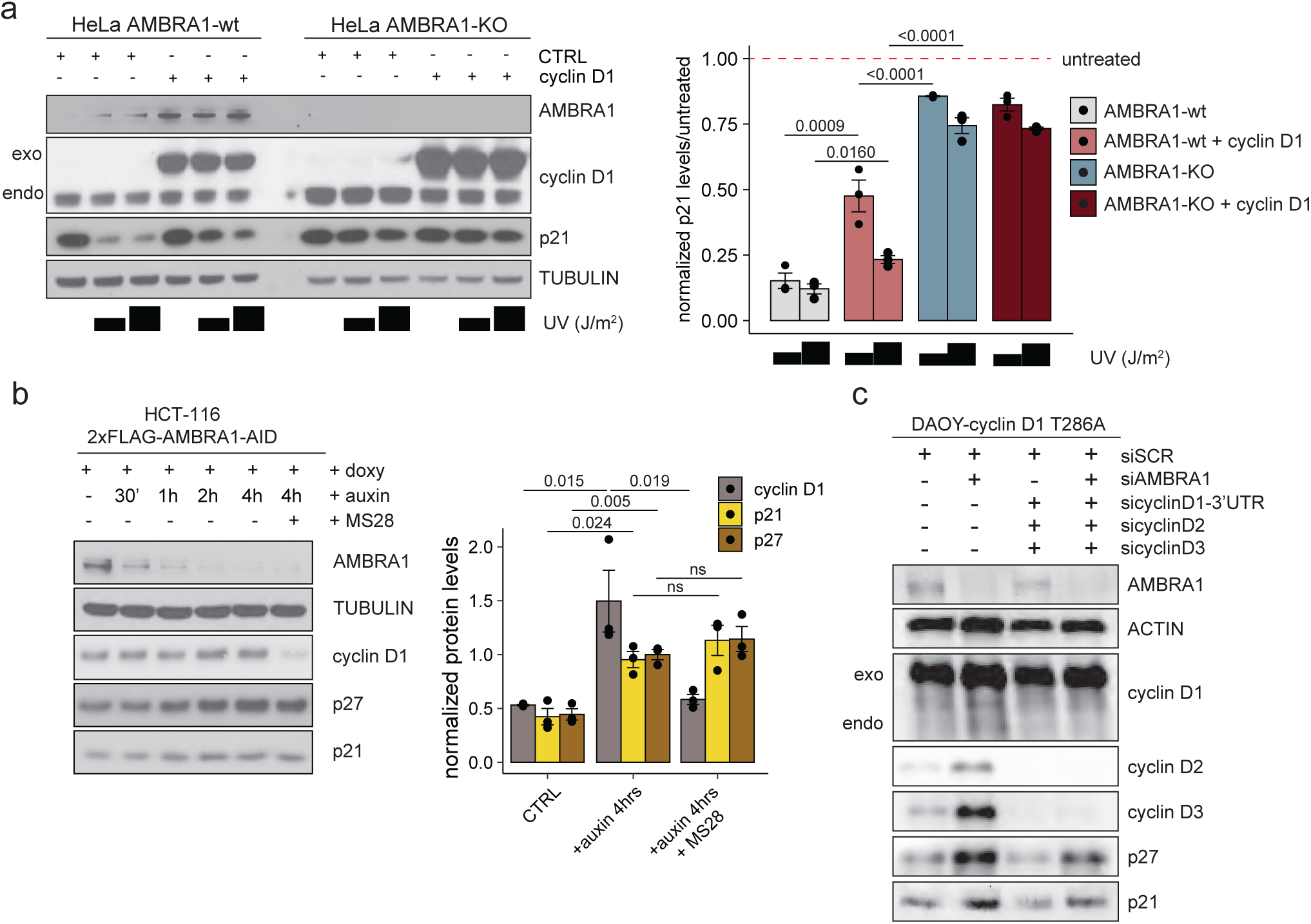
AMBRA1 mediates p21 and p27 stability both dependently and independently of cyclin D1. **a**, (*Left panel*) IB of the indicated proteins in wt or *AMBRA1*-KO HeLa cells constitutively overexpressing a doxycycline-inducible form of cyclin D1. Cells were treated with either 20 or 40 J/m^2^ UV irradiation 15’ before harvesting (n=3). (*Right Panel*) Barplot for the corresponding over CTRL-normalized quantification of p21 levels. **b**, (*Left panel*) IB for the indicated proteins of whole-cell extracts from 2×Flag-mAID-AMBRA1 HCT-116 cells pre-treated with either DMSO or doxycycline for 12 h and exposed to a combination of auxin and MS28 (cyclin D1 PROTAC degrader) for the indicated amount of time (n=3). (*Right Panel*) Barplot for the corresponding normalized quantification of p21 levels. **c**, IB for the indicated proteins in DAOY cells constitutively overexpressing the phospho-silent mutant of cyclin D1 T286A, depleted for the indicated genes with siRNA (n=3). Unless otherwise stated data are presented as mean value ±SD and n refers to biological independent samples. Data were analyzed using unpaired t-test (**a**, **b**)

### AMBRA1 regulates p21 and p27 ubiquitination via CRL4-DDB1 complex

To validate the broader impact of AMBRA1 depletion, we first confirmed in a non-MB-SHH cell line that p21 and p27 stability is affected, highlighting its widespread regulatory role by means of a CHX chase assay (Extended Data Fig. 3a). Additionally, under endogenous conditions, both p21 and p27 co-immunoprecipitated with AMBRA1 (Extended Data Fig. 3b). Multiple E3-ligases have been found to modulate cell cycle-related proteins in a time-dependent manner^26^. Among those, CRLs represent the largest E3 ubiquitin ligase family and are characterized by a modular architecture. The interchangeable cullin family members (CUL1, CUL2, CUL3, CUL4A, CUL4B, and CUL5) together with their respective adaptor proteins, such as F-box proteins or DDB1, confer substrate recognition and recruitment. To test CRLs roles in controlling the levels of cyclin D1, p27 and p21, we treated cells with MLN4924, a small molecule that inactivates CRLs by blocking cullin neddylation^27^. Like what observed upon AMBRA1 depletion in a cell-type independent manner, inhibition of CRLs resulted in the accumulation of the above three proteins (Extended Data Fig. 3c, d).

Depending on the complexes they are bound to^7,9,28^, p21 and p27 can have a dual effect in regulating G1/S transition. To identify differential protein interactors influenced by AMBRA1 loss, we immunoprecipitated p21 and p27 in parental and AMBRA1-KO cells and performed LC-MS. In the absence of AMBRA1, there was an enrichment of p21 and p27 interacting with the complex CDK4-cyclin D, suggesting an unbalanced distribution of both towards the catalytically active complexes that promote transition into the S phase (Fig. 3a, b and Extended Data Fig. 3e). The interactome shows most changes in cell cycle-related pathways, with upregulation of proteins involved in G1/S transition and p53-dependent G1 DNA damage response (Fig. 3c). Upon AMBRA1-KO, an enrichment of SKP2 among p21 interactors was observed, ruling out this E3 ligase as a potential target. This observation prompted us to investigate the specific involvement of AMBRA1 in the CRL4-DDB1 degradative pathway. To this end, p21 and p27 were immunoprecipitated from control and AMBRA1-depleted cells, followed by an assessment of DDB1’s ability to interact with both proteins. The absence of AMBRA1 impaired DDB1-p21 / -p27 interactions, suggesting AMBRA1’s direct involvement in CRL4-DDB1 ubiquitination-mediated processes (Fig. 3d). Next, we immunoprecipitated both p21 and p27, in both MB and non-MB cell lines, to assess whether the ubiquitin load of these proteins is affected by the absence of AMBRA1. The experiments showed that upon proteasome inhibition their polyubiquitination was reduced (Fig. 3e and Extended Data Fig. 3g). Since the CRL4-DDB1 complex is known to regulate p21 stability through the substrate receptor Cdt2 upon UV irradiation^3^, we investigated whether AMBRA1 functions within this axis or operates via a parallel pathway. To investigate this, Cdt2 and AMBRA1 were silenced individually or in combination in DAOY cells, and p21 levels were assessed using Quantitative Image-Based Cytometry (QIBC) and IB analysis. While depletion of either factor led to p21 upregulation, co-depletion resulted in an even greater increase in p21 levels, suggesting that AMBRA1 and Cdt2 regulate p21 through parallel mechanisms (Fig. 3f and Extended Data Fig. 3f). Finally, we performed a well-established assay to demonstrate the participation of the CRL4-DDB1 complex in the ubiquitination of p21 and p27^10^. Recombinant p21 or p27 served as substrates for the CRL4-DDB1 complex and were incubated in the presence or absence of *in vitro*–translated AMBRA1. This assay demonstrated that AMBRA1 promotes the ubiquitination of both p21 and p27, indicating its role as a positive regulator of CRL4-DDB1-mediated ubiquitination (Fig. 3g).

**Figure 3.**
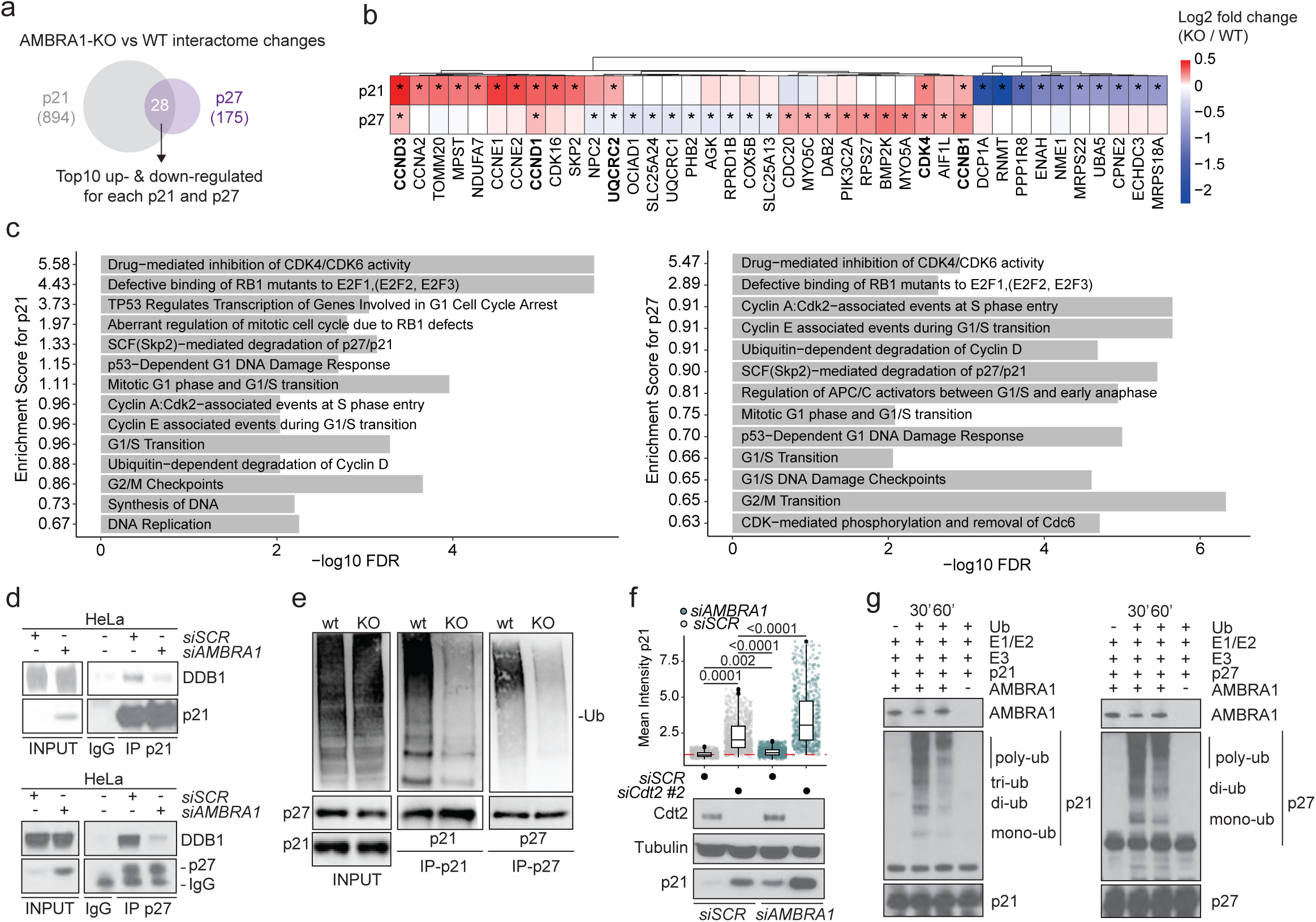
AMBRA1 mediates p21 and p27 ubiquitination. **a**, Venn diagram for shared p21 and p27 interactors in HeLa cells affected by AMBRA1 KO, obtained by LC-MS (n=4). **b**, Heatmap of top10 up- and down-regulated interactions (log2FC) from proteomics analysis in wt and *AMBRA1*-KO HeLa cells overexpressed with either p21 or p27 Flag-tagged plasmids. Asterisks for proteins indicate significant differences to wt (adjusted P value <0.05, FDR-adjusted P values from two-tailed Student’s t-test). Bold text indicates proteins shared in both interactomes. **c**, Top 14 most enriched cell cycle related biological processes (FDR < 0.01) from STRING (Functional Enrichment Analysis) affected by *AMBRA1*-KO in p21 (*Left panel*) and p27 (*Right panel*) interactomes. Bold text indicates pathways shared in both interactomes. **d**, Co-immunoprecipitation (Co-IP) of the indicated proteins (p21, *Left panel* - p27, *Right panel*) in HeLa cells depleted for AMBRA1 (n=2). **e**, IB for the *in vivo* ubiquitination levels of p21 and p27. Hela cells wt or KO for *AMBRA1* were transfected to overexpress a plasmid encoding for a HA-tagged form of ubiquitin and treated with 5 µM MG132 before harvesting as indicated (n=3). **f**, QIBC (*Top Panel*) and IB (*Bottom Panel*) analysis for the indicated proteins in DAOY cells silenced or not for the indicated genes (QIBC: n=1000 in 3 independent technical replicates; IB: n=2). **g**, *In vitro* transcribed and translated AMBRA1 and p21 or p27 were incubated in the presence of wild-type ubiquitin (Ub) and with ot without synthetic CRL4-DDB1 complex. Reactions were stopped with Laemmli buffer. Proteins were immunoblotted as indicated. Unless otherwise noted, experiments were performed at least three times. Data were analyzed using One-way ANOVA (**f**). Numerical data are available in Source Data Files.

In summary, our findings demonstrate that AMBRA1 governs the ubiquitylation and proteasomal degradation of p21 and p27, with evidence indicating that this process occurs, at least for p21, through the E3 ligase CRL4-DDB1. This mechanism operates alongside other established pathways responsible for p21 and p27 degradation, such as CRL4-DDB1^Cdt2^, CUL1-SKP2 and the one mediated by APC/C (Extended Data Fig. 3h).

### AMBRA1 regulates G1/S transition and prevents DNA damage, via cyclin D1-p21 stabilization

The initial discovery of Cip/Kip family members characterized them as nuclear proteins with a predominant function of inhibiting cyclin–CDK activity and -hence-, cell-cycle progression^29^. However, building evidence has clarified their dual role: they slow the cell cycle by inhibiting the CDK2-cyclin E/A complex while simultaneously promoting the G1/S transition through the stabilization of the CDK4-cyclin D complex^7^. To explore in depth the role of AMBRA1 in the regulation of cyclin D1-p27/p21 and the G1/S transition, we depleted AMBRA1 in several cell types, including our MB cell models, and quantified the proportion of cells at different cell cycle phases. Consistently with previous findings, the absence of AMBRA1 resulted in the accumulation of cells in the S phase with the concomitant accumulation of DNA damage, only in an Rb-proficient genetic background^30^ (Fig. 4a and Extended Data Fig. 4a, b). To further validate the role of Rb and the accumulation of S phase cells in the RS phenotype observed in AMBRA1-deficient cells, we co-depleted AMBRA1 and Rb in ONS76 cells. QIBC analysis demonstrated that Rb depletion alone increased the number of cells in S phase and was associated with an increase in γH2AX signal (Extended Data Fig. 4c). Notably, this increase was not readily detectable by IB, except upon additional CHK1 inhibition, (Extended Data Fig. 4d), likely reflecting the greater sensitivity of the QIBC single-cell assay for capturing changes occurring in specific subpopulations. Although co-silencing of AMBRA1 and Rb did not further increase the number of cells in S phase, AMBRA1 depletion further enhanced γH2AX levels and CHK1 activation. (Extended Data Fig. 4c-d). This suggests that the RS induced by AMBRA1 depletion arises from at least one other source besides accelerated entry into S phase. To quantitatively evaluate the molecular dynamics occurring among cyclin–CDK in the absence of AMBRA1, we performed Proximity Ligation Assay (PLA) and found that hyper-stabilization of CDK4-p21/p27 caused by AMBRA1 depletion prevents Cip/Kip proteins to associate with CDK2 (Fig. 4b). Using a reporter system to measure CDK4/6 and CDK2 activities in individual cells^31^, increased activity was observed in both basal and serum-deprived conditions, driving S-phase entry independently of mitogenic stimuli (Extended Data Fig 4e, f). Recent insights into CDK4/6–CDK2 dynamics suggested that the CDK4/6 inhibitor palbociclib inhibits only cyclin D–CDK4/6 dimers, but not trimeric cyclin D–CDK4/6-p27. In the first case, palbociclib might prevent the formation of active CDK4-containing complexes (through binding to CDK4), and indirectly block CDK2 by liberating Cip/Kip inhibitors^32^. Additionally, it was found that the FDA-approved CDK4/6 inhibitors have different target spectra, with palbociclib mainly inhibiting CDK4/6, while abemaciclib also influencing CDK2 and CDK1^33^. Based on that, we hypothesized that, despite being palbociclib-resistant^24^, AMBRA1-depleted cells would retain sensitivity to abemaciclib. Of note, the experimental confirmation came from QIBC analysis, which showed that abemaciclib, unlike palbociclib, was able to fully rescue the S-phase shift and hold cells in G1 by restricting both CDK4/6 and CDK2 activity in AMBRA1-deficient cells (Extended Data Fig 4g-i).

**Figure 4.**
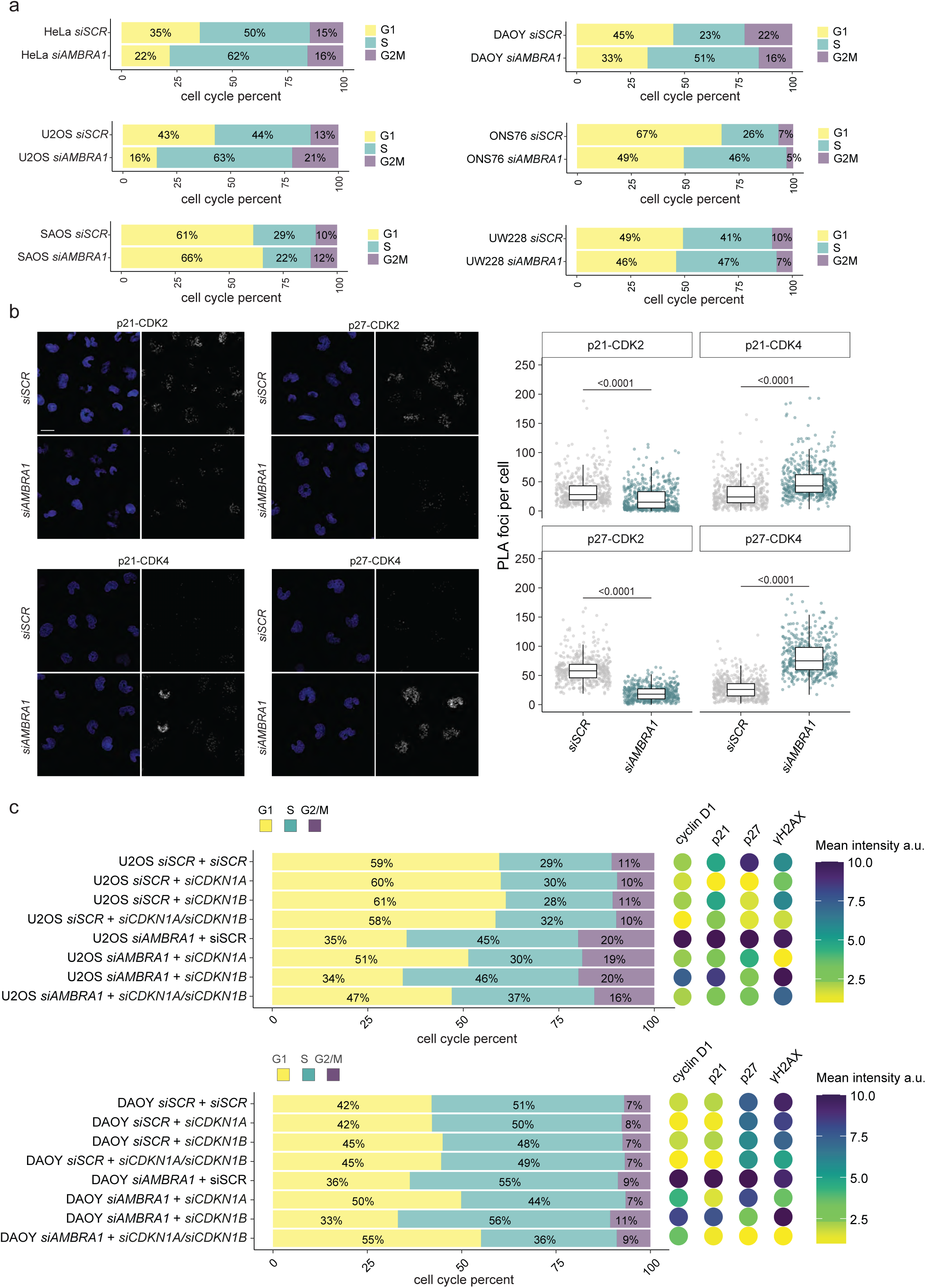
AMBRA1 regulates G1/S phase transition and DNA damage via cyclin D1-p21 stabilization. **a**, Stacked barplot of cell cycle distribution (%) of QIBC analysis for a scramble or AMBRA1-depleted HeLa (*Top Left panel*), U2OS (*Middle Left panel*), SAOS (*Bottom Left panel*), stained for EdU and Hoechst (at least 2000 cells were acquired per condition). Stacked barplot of cell cycle distribution (%) of FACS analysis for scramble or AMBRA1-depleted DAOY (*Top Left panel*), ONS76 (*Middle Left panel*), UW228 (*Bottom Left panel*), expressing FUCCI-CA reporter (at least 10000 cells were analyzed per condition), n=3. **b**, *Left panel*, Confocal imaging of PLA for the indicated proteins, in control or AMBRA1-depleted DAOY cells. *Right Panel*, Boxplot (median values with IQR) of foci number per cell, for each separate PLA reaction (n=1000 from 5 independent biological replicates). Scale bar = 50 um. **c**, Stacked bar plot of QIBC analysis for normalized cell cycle phases distribution (%) in U2OS (*Top panel*) and DAOY (*Bottom panel*) cells depleted for the indicated genes with siRNA, with the respective bubble plot showing normalized mean intensity (a.u.) of the indicated proteins (cyclin D1: n=1000 / p21: n=1000 / p27: n= 1000 / γH2AX: n=1000 in three independent technical replicates). To separate the different cell cycle phases cells were stained for EdU and Hoechst. Unless otherwise stated data are presented as mean value and n refers to technical independent samples. Data were analyzed using a two-tailed unpaired t-test (**b**).

To gain insight into the effect of hyper-stabilization of p27 and p21 in the CDK4-cyclin D1-p21/p27 holoenzyme, we depleted p21 and p27 individually or in combination with AMBRA1 in U2OS and DAOY cells and measured the cell cycle distribution, and the protein expression of cyclin D1, p21, p27 and γH2AX. Intriguingly, the cell cycle shift and the DNA damage caused by AMBRA1 depletion could be restored by p21 interference but not p27, suggesting an epistatic role of p21 in licensing premature S-phase entry (Fig. 4c). This may force cells to synthesize their DNA using a limited number of origins of replication^34^. However, the absence of AMBRA1 showed no reduction in phospho-MCM2, a surrogate marker of origin activation^35^, suggesting no significant changes in their number (Extended Data Fig. 4j). Together, these results imply that the RS resulting from AMBRA1 depletion stems from disrupted proteostasis of the cyclin D-p21 complex with direct negative consequences for the replication forks.

### AMBRA1 regulates replication fork speed and lagging strand synthesis via p21-PCNA dynamics

In the absence of AMBRA1, cells experienced increased RS presumably due to the enhanced stability of cyclin D1-3^2^, however, the mechanism of how RS occurs is still elusive. As previously observed in other cell types^2^, in the absence of AMBRA1, MB cell lines showed an accelerated speed of fork progression with the accumulation of asymmetric forks, a readout of compromised fork stability (Fig. 5a and Extended Data Fig. 5a). Next, we tested whether the acceleration of forks could leave under-replicated DNA and performed the S1-nuclease assay to reveal ssDNA gaps^36^. Indeed, in the absence of AMBRA1, DNA replication forks were sensitive to the S1 digestion (Fig. 5b). Higher replication fork speed may result from reduced origin firing. When fewer replication origins are activated, existing replication forks compensate by increasing their speed to ensure timely DNA replication^37^. To determine whether the effects observed upon AMBRA1 deficiency were due to a direct impact on the replisome or an indirect consequence, we measured inter-origin distance (IOD) both under basal conditions and following ATR inhibitor (ATRi) treatment, which promotes the activation of dormant origins^38,39^. Notably, no differences in IOD between AMBRA1-depleted and control cells were found (Fig. 5c). To further investigate the early steps of replication initiation, we measured chromatin-bound levels of key origin licensing and activation proteins, namely MCM2, CDC6, CDC45, and CDT1, in both MB and non-MB cell lines (Extended Data Fig. 5b-g). In line with the previous analysis, no consistent discrepancies were observed in AMBRA1-depleted cells across the three cell lines, suggesting that the replication stress induced by AMBRA1 deficiency may occur independently of origin initiation. In agreement with the altered processivity of the replisome in DNA replication and the presence of under-replicated DNA, AMBRA1 silencing profoundly impacted S-phase dynamics, as confirmed by live-imaging analysis. While S-phase entry and its initial steps were accelerated, the late S-phase and transition into G2 were prolonged, suggesting delays in traversing the S-G2 checkpoint. These findings are in line with our previous observations in non-cancerous neural stem cells^2^. Nevertheless, despite these delays, the overall duration of the cell cycle remained shortened in AMBRA1-silenced cells (Extended Data Fig. 5h).

**Figure 5.**
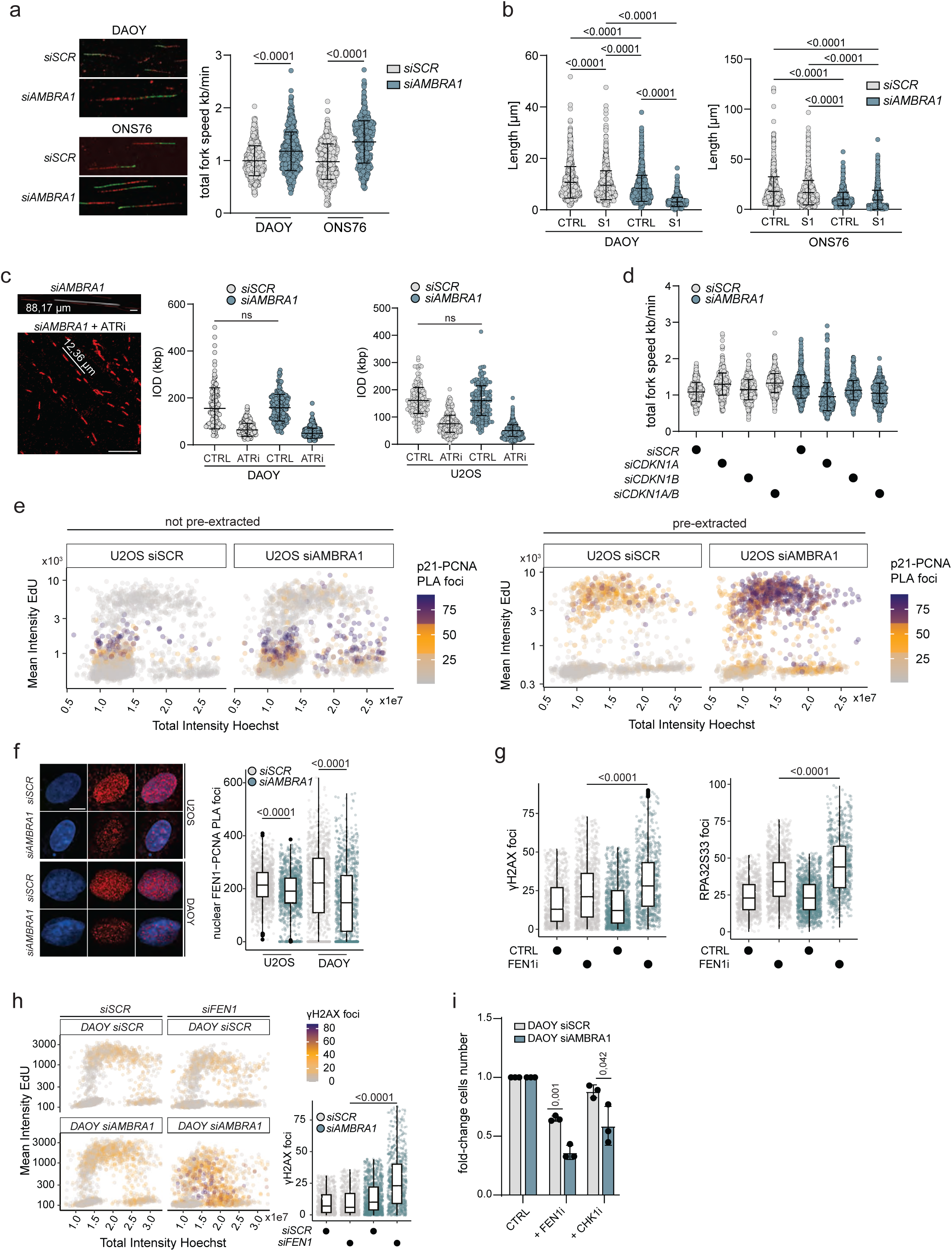
AMBRA1 maintains replication fidelity by modulating p21-PCNA interactions and lagging strand maturation. **a**, Replication fork speed from DAOY and ONS76 cells depleted or not for AMBRA1 (DAOY *siSCR*: n = 546; DAOY *siAMBRA1*: n = 537 / ONS76 *siSCR*: n =576; ONS76 *siAMBRA1*: n = 552; in three technical independent experiments). **b**, Replication fork length in DAOY and ONS76 cells depleted or not for AMBRA1 and treated without (CTRL) or with S1 nuclease (S1) for 30’ (DAOY *siSCR* CTRL: n = 1002; DAOY *siSCR* + S1: n = 1008; DAOY *siAMBRA1* CTRL: n = 1002; DAOY *siAMBRA1* + S1: n = 1005 / ONS76 *siSCR* CTRL: n =1010; ONS76 *siSCR* + S1: n = 623; ONS76 *siAMBRA1* CTRL: n = 764; ONS76 *siAMBRA1* + S1: n = 1010; in three technical independent experiments). **c**, (*Left panel*) Representative images of Replication Inter-Origin Distance (IOD) analysis and the associated quantification in DAOY and (*Middle panel*) U2OS (*Right panel*) cells depleted or not for AMBRA1 and treated without (CTRL) or with ATRi for 30’ (DAOY *siSCR* CTRL: n = 126; DAOY *siSCR* + ATRi: n = 229; DAOY *siAMBRA1* CTRL: n = 148; DAOY *siAMBRA1* + ATRi: n = 368 / U2OS *siSCR* CTRL: n =162; U2OS *siSCR* + ATRi: n = 296; U2OS *siAMBRA1* CTRL: n = 177; U2OS *siAMBRA1* + ATRi: n = 398; in three technical independent experiments). **d**, Replication fork length in U2OS cells treated as in **4c** (U2OS *siSCR* + *siSCR*: n = 513; U2OS *siSCR* + *siCDKN1A*: n = 561; U2OS *siSCR* + *siCDKN1B*: n = 516; U2OS *siSCR* + *siCDKN1A*/*B*: n = 564; U2OS *siAMBRA1* + *siSCR*: n =533; U2OS *siAMBRA1* + *siCDKN1A*: n = 527; U2OS *siAMBRA1* + *siCDKN1B*: n = 540; U2OS *siAMBRA1* + *siCDKN1A*/B: n = 564; in three technical independent experiments). **e,** QIBC analysis of PLA reaction for p21 and PCNA in scramble or AMBRA1-depleted U2OS. Before fixation cells were either pre-extracted (*Right panel*) or not (*Left panel*), then PLA reaction for p21-PCNA was performed and lastly cells were stained for EdU and Hoechst (2000 cells are displayed per the condition of three independent replicates). **f**, Representative images of QIBC analysis of PLA reaction for PCNA and FEN1 in scramble or AMBRA1-depleted U2OS and DAOY cells with its corresponding jittered boxplot quantification (n=1000 per condition in 3 independent biological replicates). Scale bar = 10 um. **g**, Jittered boxplot quantification of QIBC analysis of γH2AX (*Left panel*) and phospho-RPS32(S33) (*Right panel*) foci in scramble or AMBRA1-depleted DAOY S-phase cells treated or not with FEN1 inhibitor (n=1000 per condition in 3 independent biological replicates). **h**, (*Left panel*) QIBC analysis of γH2AX foci along cell cycle in DAOY cells depleted for the indicated genes stained for EdU and Hoechst (2000 cells are displayed per condition). (*Right panel*), Jittered boxplot quantification of γH2AX foci (n=1000 per condition in 3 independent biological replicates). **i**, Barplot representing fold-change cell number in DAOY cell line depleted or not for AMBRA1, treated for 48 hrs with the indicated cytotoxic compounds (n=3). Data are presented as mean value ±SD for scatter plots while median and IQR is shown for boxplots. n refers to biological independent samples. Data were analyzed using One-way ANOVA (**a**, **b**, **c, d, f, g, h**).

To further support the observation that p21 silencing rescues cell cycle alterations and to establish a causal link between the AMBRA1 phenotype and p21’s role, we measured replication fork speed in U2OS cells after depleting p21 and p27 individually or in combination with AMBRA1. Consistent with our previous observation, p21 depletion accelerated fork speed^40^, in contrast to p27. Furthermore, p21 knockdown had a predominant effect in reducing fork speed acceleration caused by AMBRA1 silencing (Fig. 5d and Extended Data Fig. 5i). Because it is known that among all biological PIP-boxes p21 has the highest known affinity for PCNA^41^, we tested whether this interaction is altered in the absence of AMBRA1, taking advantage of a high-content microscopy method to quantify the signal of PLA foci in chromatin-unbound and -bound p21-PCNA complex. The results showed that upon AMBRA1 silencing the interaction is significantly enhanced in distinct cell cycle phases (Fig. 5e and Extended Data Fig. 6a). Notably, when analyzing the soluble fraction, the enhancement was prominent during the G1/S transition. However, analysis of the chromatin-bound fraction revealed significant enrichment throughout the S phase, suggesting a potential impact on PCNA regulation. This effect is likely driven by a general accumulation of p21 across all phases of the cell cycle in the absence of AMBRA1 (Extended Data Fig. 6b), which may exacerbate the disruption of replication-associated protein interactions.

FEN1, a 5’-3’ exonuclease essential for Okazaki fragment maturation, cannot simultaneously bind PCNA with p21, as the formation of a p21-PCNA-FEN1 complex is precluded *in vitro*^42^. Moreover, p21 overexpression disrupts the FEN1-PCNA interaction and reduced FEN1 levels are associated with accelerated replication fork progression^40^. Based on these findings, we hypothesized that the hyper-stabilization of p21 might impair lagging strand synthesis by altering the FEN1-PCNA interaction. To test this, PLA analysis of nuclear FEN1-PCNA interactions using QIBC was performed and revealed a significant decrease in their association upon AMBRA1 depletion (Fig. 5f). In parallel, QIBC analysis revealed that enhancing the impact on lagging-strand synthesis through FEN1 inhibition or silencing increased DNA damage. This was reflected in higher γH2AX levels, enhanced RPA32 (S33) foci formation, and significant alterations in cell cycle distribution (Fig. 5g, h). To assess the functional consequences of these effects, we performed cell viability assays in AMBRA1-deficient cells. As a positive control, we included CHK1/2 inhibitor AZD7762, which we previously described to induce synthetic lethality when combined with AMBRA1 depletion². Accordingly, low AMBRA1 expression significantly sensitized DAOY cells to CHK1, or FEN1 inhibitor, whereas no such effect was observed in ONS76 cells, which already exhibited a drastic decrease in cell viability upon AMBRA1 depletion alone (Fig. 5i and Extended Data Fig. 6c, d).

These results collectively highlight the essential role of AMBRA1 in ensuring proper DNA replication by regulating p21 dynamics at the replisome.

### *TP53* status dictates sensitivity to AMBRA1 depletion and RS in MB-SHH

While both MB cell lines exhibited a similar pattern of RS induction, only the ONS76 displayed enhanced DNA damage which led to an increase in the apoptotic fraction (Fig. 6a), suggesting that these cells are unable to cope with high levels of RS. Notably, the observed molecular and phenotypic differences among these cell lines may stem from variations in their mutational profiles. Interestingly, while ONS76 retains a functional *TP53* (as reported in cellosaurus.org), both DAOY and UW228 harbor point mutations in the tumor suppressor (DAOY: G725T / UW228: C464A)^43^. In line with this, DepMap Gene Effect analysis revealed that TP53 KO had the most significant positive effect on cell viability in the ONS76 (Fig. 6b). To determine whether the detrimental effects of AMBRA1 silencing on ONS76 cells are linked to p53 activity, we engineered these cells for stable p53 downregulation, replicating the conditions observed in the other two MB cell lines. Importantly, IB analysis showed that p53 knockdown could indeed mitigate γH2AX levels as well as PARP cleavage boost caused by AMBRA1 depletion (Fig. 6c, d). Accordingly, the absence of p53 rescued the sharp decline in cell viability, restoring it to control levels (Fig. 6e). Further supporting the genetic interplay between *AMBRA1* and *TP53*, DepMap Gene Effect analysis revealed a strong co-dependency between the two genes, with MB cell lines predominantly aligning along the regression line. Notably, p53 emerged as the sixth most co-dependent gene with AMBRA1 (Fig. 6f). Finally, reinforcing this relationship across different MB-SHH subtypes, AMBRA1 mRNA levels were the lowest in the subtype SHH-α, which is characterized by the highest prevalence of *TP53* mutations^44^ (Fig. 6g).

**Figure 6.**
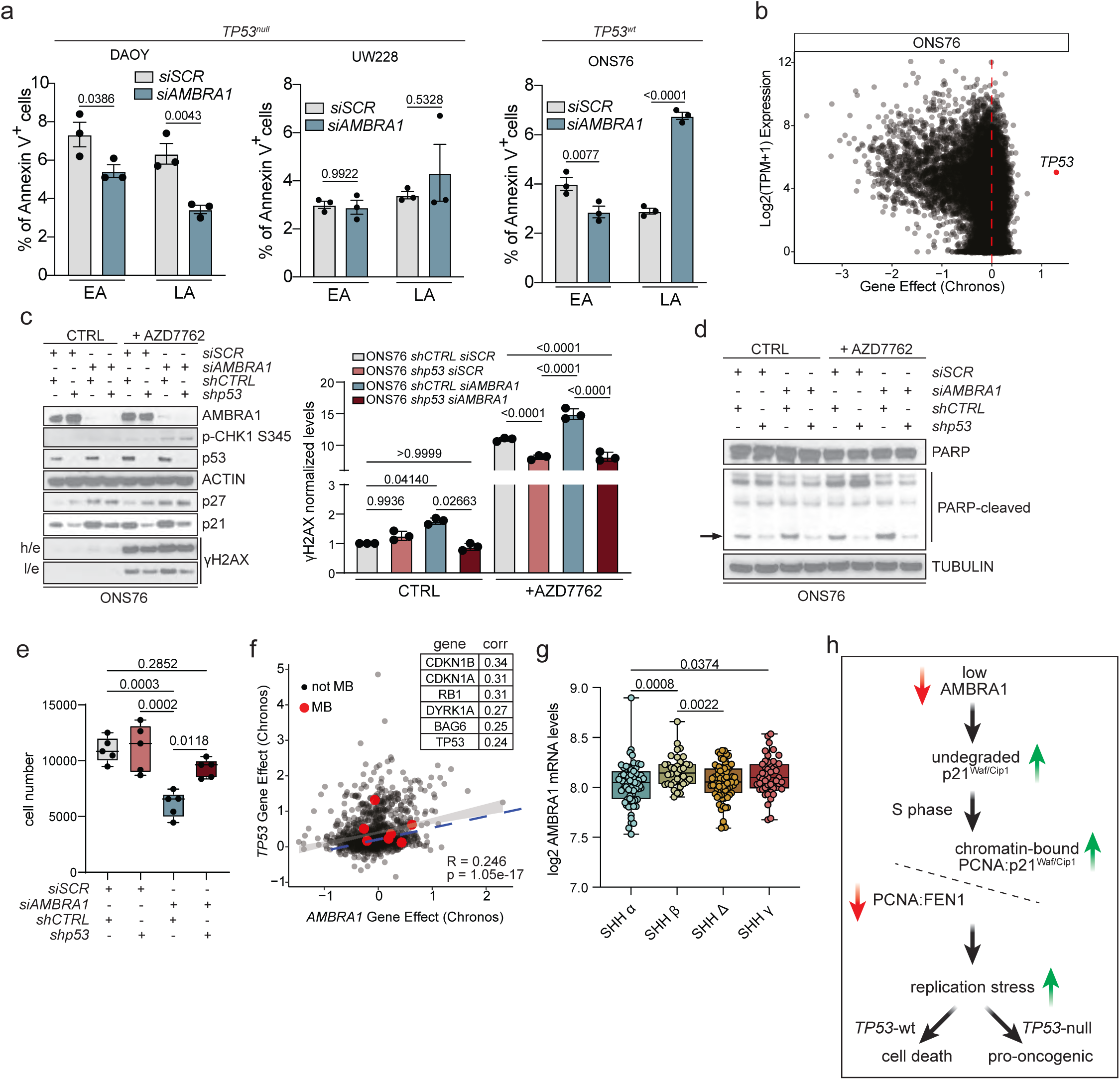
*AMBRA1-TP53* dependency links RS response to cell viability in MB. **a,** Barplot of Annexin V^+^ cells (%) in early and late apoptotic phase (EA - LA) detected by flow cytometry with Annexin V-APC staining and PI counterstain, in DAOY, UW228 and ONS76 in control or AMBRA1-depleted condition (n=3). **b**, Scatter plot distribution of Chronos corrected GeneEffect of CRISPR KO over mRNA expression in ONS76 cell line from DepMap. **c**, (*Left panel*) IB for the indicated proteins of whole-cell extracts from control (*shCTRL*) or p53 constitutively downregulated ONS76 cells in scramble or AMBRA1-depleted (*shp53*). 24 hrs before harvesting cells were treated without or with 200 nM AZD7762 (n=3). (*Right panel*) Barplot of γH2AX normalized protein levels observed in **c** (n=3). **d**, IB for the indicated proteins of whole-cell extracts from control (shCTRL) or p53 constitutively downregulated ONS76 cells in scramble or AMBRA1-depleted. 24 hrs before harvesting cells were treated with or without 200 nM AZD7762 (n=3)**. e**, Boxplot of cell number from shCTRL or *shp53* ONS76 cells in a scramble or AMBRA1-depleted condition. **f**, (*Left panel*) Scatter plot distribution of Chronos corrected GeneEffect of CRISPR KO for *AMBRA1* and *TP53* from 24Q4 DepMap release; MB cell lines in red. (*Right panel*) Table with Top6 co-dependent gene with AMBRA1. **g**, Boxplot for mRNA log2 expression of AMBRA1 in the different SHH subgroups derived from the publicly available dataset Cavalli (763 patients, fpkm normalized, mb500rs1 chip). SHH-α = 65; SHH-β = 35; SHH-Δ = 76; SHH-γ = 47. **h**, A comprehensive model illustrating how reduced AMBRA1 levels lead to the accumulation of RS. Data were analyzed using either with two-way ANOVA (**a**), or one-way ANOVA (**b**, **e**) Welch’s two-tailed unpaired t-test (**g**)

## Discussion

Despite decades of intense research on Cip/Kip protein families, p21 and p27 are still often considered as mere cell cycle inhibitors^45–48^, however, multiple evidence including the present manuscript expands the perception of these two key regulatory components of the CDK4/6 kinase complex^49^. In particular, our data provides insight into the coordinated degradation pathways of p21 and D-type cyclins, highlighting a key role for the substrate receptor AMBRA1 in mediating their turnover via the CRL4-DDB1 complex. This finely tuned regulation occurs throughout the cell cycle and is critical for maintaining the appropriate stoichiometry and function of the CDK4/6–cyclin D–p21 complex, which in turn supports proper S-phase entry and safeguards DNA replication fidelity (Extended Data Fig. 7a). Importantly, our findings add to a growing appreciation of the nuanced, context-dependent interplay between p21, p27, Cyclin D1, and their associated complexes. These proteins can mutually regulate each other’s stability and function, resulting in complex network dynamics that are sensitive to cell type, signaling cues, and genetic background. Our results reinforce the view that dysregulation of any single component can propagate through the network to influence cell cycle progression and replication stress responses.

Indeed, beyond its well-established role in regulating the G1 checkpoint, p21 also coordinates DNA replication dynamics and influences the cellular response to replication stress^50,51^. Previously, in an Rb-null model of senescence evasion, we demonstrated that chronic overexpression of p21 negatively impacts the speed of replication forks^52^. Conversely, we also observed an opposite effect upon p21 depletion where the replication fork speed is enhanced in a PARP1-dependent manner^40^. Here, we enrich the broad spectrum of p21 involvement in replisome processivity by mechanistically defining how its acute stabilization, combined with the accelerated transition from G1 to S phase, caused by AMBRA1 silencing, affects the dynamic interaction between PCNA and its partners. Dysregulated levels of p21 impact negatively the lagging strand DNA synthesis, enhancing RS and making AMBRA1-depleted cells sensitive to FEN1 inhibitors (Fig. 6h). Our results suggest that p21 levels are tightly regulated during S phase to ensure proper DNA replication. Notably, AMBRA1 depletion disrupts this balance, leading to aberrant regulation of p21. Interestingly, both p21 depletion and enhanced stabilization of the protein result in accelerated replication fork progression. This phenomenon is likely attributable to dysregulated synthesis of the lagging DNA strand in both scenarios.

In our attempt to find a specific tumor context highly relying onto this aberrant modulation, we identified a defined subset of MBs, namely MB-SHH, where we demonstrate the importance of AMBRA1 downregulation in enabling tumor cells proliferation independently of mitogenic signaling. Our results also document the relevance of the G1/S regulatory function of AMBRA1 in cerebellar development, potentially shedding new light on the involvement of this molecular axis in other cerebellar-related disorders. Additionally, by using genetic, biochemical and pharmacological approaches, we mechanistically define how AMBRA1 depletion concomitantly exacerbates CDK4/6 as well as CDK2 activity, leading to palbociclib resistance in MB-SHH cell lines. Nonetheless, we identify an alternative CDK4/6 inhibitor, abemaciclib, as a promising therapeutical approach, given the significant repressive effect this compound has on CDK2 as well^33^.

Of clinical relevance, previous observation from a SB transposon-based screening found AMBRA1 as one of the most frequently mutated genes in a cohort of embryonal tumors of the nervous system primed by p53 mutation^17^. In agreement with that, we describe how p53 mutation supports MB-SHH cells in coping with the intrinsic genomic instability caused by AMBRA1 deficiency, licensing aberrant S-phase entry and promoting apoptotic evasion. While our data from both cell lines and tumor samples support a potential role for AMBRA1 in SHH-MB, further studies incorporating clinically relevant models and larger patient cohorts will be essential to conclusively define its contribution to SHH-MB initiation and progression *in vivo*. In summary, our findings establish AMBRA1 as a critical regulator of S-phase entry and DNA replication fidelity through its dual role in controlling p21 turnover and cyclin D expression. Depletion of AMBRA1 leads to aberrant stabilization of the CDK4/6–cyclin D–p21/p27 complex, accelerated G1/S transition, and impaired coordination of lagging strand synthesis, ultimately promoting replication stress and genomic instability. Our work not only uncovers a mechanistic basis for AMBRA1’s tumor suppressor function in SHH-MB, but also highlights abemaciclib and FEN1 inhibition as a potential therapeutic strategy to counteract the resistance conferred by AMBRA1 deficiency. These insights have direct implications for the development of targeted therapies and biomarker-driven patient stratification in medulloblastoma and potentially other AMBRA1-deficient cancers.

## Acknowledgements

G.M. is/has been supported by AIRC Fellowship (Ref. 26727), by the Danish Cancer Society (R302-A17590), by Horizon-MSCA-2022 Postdoctoral Fellowship (Project 101108184). F. Cecconi labs are supported by AIRC IG23543, Novo Nordisk 0070834, KBVU R231-A14034 and R325-A19075, PRIN 2017FS5SHL and 2020PKLEPN from the Italian Ministry of Research, RF-2019-12369700 from the Italian Ministry of Health, and Danmarks Grundforskningsfond (DNRF125). J.B. laboratory is supported by grants from The Danish Council for Independent Research (number DFF-1026-00241B), The Novo Nordisk Foundation (grant number 0060590), The Swedish Research Council VR-MH 2014-46602-117891-30 and The Swedish Cancer Foundation/Cancerfonden (number 170176). A.M.-M. laboratory is supported by grants, KBVU R302-A17590. F.C., J. B. and A. M.-M. laboratories in Copenhagen are part of the Center of Excellence for Autophagy, Recycling and Disease (CARD), funded by the Danmarks Grundforskningsfond (DNRF125). VC is supported by the Italian Ministry of Health (GR-2021-12372771), by the MIUR (PRIN 2022 Bando PNRR P2022TJZYZ to VC; MIUR-Italy Departments of Excellence 2023-2027 to the Dept. of Science, Univ. Roma Tre). FN’s lab is supported by the “Associazione Italiana Ricerca sul Cancro” (My First Airc Grant -Ref. 27019), by “CureSearch Young Investigator Awards in Pediatric Oncology Drug Development”, by “Fondazione Roche per la Ricerca Indipendente”, by the Italian Ministry of Health (GR-2019–12369231), by the Italian Ministry of University and Research (PRIN PNRR P2022JLHZZ). G.C. is enrolled in the PhD Program in Cellular and Molecular Biology, Department of Biology, University of Rome “Tor Vergata. We give a special thanks to C. Dinant and the BioImaging Core Facility at the Danish Cancer Institute, to Plaisant S.r.l. (Castel Romano) animal facility for their invaluable help with the in vivo experiments, and Dr. Carlo Rodolfo (Rome) for his strenuous help and support.

## Author contributions

G.M., A.M-M, and F.C. conceived the study and designed experiments. G.M., G.C., A.A.R, C.F. carried out the biochemical and microscopy experiments linking AMBRA1 to p21 and p27 regulation, genomic instability, and synthetic lethality. G.M and C.D.S performed analyses regarding Ambra1 cKO cerebellar development. K.K. carried out the S1 assay. P.G. performed LC/MS analysis and S.M. carried out of data visualization and statistical analysis for mass spectrometry data. D.Q., S.R., A.M., F.L. provided the medulloblastoma patients samples and performed the IHC scoring. F.N., V.C., J.B., provided critical support, key data analyses, and conceptual advice. G.M., A. M.-M. and F.C. wrote the original draft. All authors took part in writing, reviewing, and editing the final manuscript. All authors read and accepted the manuscript.

## Materials & Correspondence

Correspondence to:

Francesco Cecconi

Apolinar Maya-Mendoza

## Methods

### Statistics and reproducibility

Statistical analyses were performed using GraphPad Prism 9 or R (v4.2.1) software. The statistical tests used are indicated in the figure legends.

### Generation of Ambra1^flox/flox^/Nestin-cre mice

To generate Ambra1^flox/flox^/Nestin-cre^2^ homozygous Ambra1^flox/flox^ females were bred onto Ambra1^wt/flox^/Nestin-cre. Nestin-cre were purchased from Jackson Lab and maintained in the animal facility Plaisant Castel Romano (RM, Italy). Mutants were genotyped by PCR of genomic DNA extracted from tails or ears using the following primers:

*Tm1c* FW 5’-TGATAGTCCACGCTCGACCT-3’,

*Tm1c* RV 5’-CTAATCCGCCTACTGCGACT-3’,

*Ambra1* wt FW 5’-TCTGGTTGCCTAGATGGGGA-3’,

A*mbra1* wt RV 5’-ACTCATGTTAGAGCCTCCTGC-3’,

*Cre* FW 5’-CGGTCGATGCAACGAGTGATGAGG-3’,

*Cre* RV 5’-CCAGAGACGGAAATCCATCGCTCG-3’,

All in vivo experiments were approved by and performed following the ethical international, EU and national requirements and were approved by the Italian Health Ministry (D.lgs 26/2014; N°. 737 03/2013; N°88/2016-PR).

### Cell culture

All cell lines were grown at 37 °C in a humidified incubator containing 5% CO2.

U2OS, HeLa, SAOS and cells were purchased from the American Type Culture Collection (ATCC) and cultured in Dulbecco’s modified eagle’s medium (DMEM) GlutaMAX^TM^ (Gibco) supplemented with 10 % fetal bovine serum (FBS) (Gibco) and antibiotics. DAOY were a kind gift of Dr. A. Di Giannatale, and were cultured in MEM Medium (Gibco), supplemented with 10% FBS (Gibco) and antibiotics. ONS76 cells were purchased from Accegen and cultured in RPMI-1640 Medium (Gibco), supplemented with 10% FBS (Gibco) and antibiotics. UW228 were a kind gift of Dr. D. Raleigh, and were cultured in DMEM Medium (Gibco), supplemented with 10% FBS (Gibco) and antibiotics. HCT-116 2×Flag-mAID-AMBRA1 cells stably infected with pTRIPZ-HA-TIR1 were a kind gift of Prof. M. Pagano and were maintained in McCoy’s 5A medium supplemented with 10% Tet System Approved FBS (Takara, Clontech Laboratories) and 1% penicillin/streptomycin/l-glutamine (Corning Life Sciences).

Monoclonal DAOY-FUCCI, ONS76-FUCCI and UW228-FUCCI cells were generated as previously described^53^ and maintained in the same condition as their parental progenitors. Monoclonal DAOY-DHB-mVenus-p2a-mCherry-CDK4KTR, ONS76-DHB-mVenus-p2a-mCherry-CDK4KTR cells were generated as previously described^31^ and maintained in the same condition as their parental progenitors. Hela wt and KO for AMBRA1 were generated previously^54^ and were cultured in DMEM Medium (Gibco), supplemented with 10% FBS (Gibco) and antibiotics.

### Transfections, plasmids and siRNAs

Transient overexpression was carried out using Lipofectamine 2000 or Lipofectamine 3000, depending on the experimental conditions and according to the manufacturer instructions (Invitrogen). Plasmids for transient overexpression coding for AMBRA1 wt and AMBRA1-myc were constructed as previously described^5^. The HA-UBIQUITIN plasmid was used for transient overexpression of ubiquitin in HeLa cell line^55^. *Homo Sapiens* cyclin D1-FLAG and cyclin D1-T286A-FLAG cDNAs were generated as in^2^. For stable overexpression of cyclin D1/D2/D3 lentiviral pLVX-IRES-mCherry lentiviral vectors containing N-terminal Flag or Strep-tags were used, and pLVX-TREGS-GFP-3xFlag plasmid was used as a negative control. FFSS indicates a tandem 2×Flag–2×Strep tag.

For stable overexpression of cyclin D1 in the HeLa cell line, lentiviral pTripz-FFSS-CCND1 was used and pLVX-GFP-3xFlag plasmid was used as negative control. To generate different stable FUCCI CA (5) cell lines the following plasmids were used in a 1:4 ratio: piggyBac-CMV-HA-tag PBase (a gift from Tiberi lab) and FUCCI CA (5) (Addgene #153521^53^, gift from Atsushi Miyawaki). The following plasmids were used to generate the ONS76 shCTR and shp53 stable cell line: pLKO.1-puro – CMV - TurboG (Sigma-Aldrich) and pLKO.1 - puro - shp53^56^ (Addgene #19119), a gift from Bob Weinberg.

siRNA transfections were carried out using Lipofectamine RNAiMAX (Invitrogen) as described by the manufacturer. RNA interference was performed using the oligonucleotides as previously described^7^. MISSION® siRNA Universal Negative Control #1 (Sigma) was used as siRNA control. Employed siRNA sequences are as follows:

*siAMBRA1*-5’UTR: 5’-GGACAACUUACAAGGACCU-3’

*siAMBRA1*: 5’-GAGUAGAACUGCCGGAUAG-3’;

*siCCND1 – 3’UTR*: 5’-GCGUGUAGCUAUGGAAGUU-3’

*siCCND2:* 5’-GAUCGCAACUGGAAGUGUG-3’

*siCCND3:* 5’-UGCGGAAGAUGCUGGCUUA -3’

*siCDKN1A:* 5’-CUGUCACAGGCGGUUAUGA-3’

*siCDKN1B:* 5’-ACGUAAACAGCUCGAAUUA-3’

*siCDKN1A-3’UTR:* 5’-CGCTCTACATCTTCTGCCTTA-3’

*siCDKN1B-3’UTR:* 5’-GTAGGATAAGTGAAATGGATA-3’

*siDDB1*: 5’-GCUGAGUGCUUGACAUACCUUGAUA-3’

*siSKP2:* 5’ *-*GCUGCGCAUCCGCAGUAGUU-3’

*siFEN1*: 5’ *-*GATGCCTCTATGAGCATTTAT-3’

*siCDT2 #1*: 5’ *-*CTGGTGAACTTAAACTTGTTA-3’

*siCDT2 #2*: 5’ *-*GCTCCCAATATGGAACATGTA-3’

### Virus production and infection

For lentiviral production vectors were produced by transfecting human embryonic kidney (HEK) 293T cells with the Fugene (Promega) in accordance with the manufacturer’s instructions. After removal of cell debris with filter, lentiviruses contained in the supernatant were precipitated with Lenti-X concentrator (TakaraBio Cat. No. 631231) and resuspended in PBS. Depending on the experimental condition cells were plated at low confluency (20-30%) and then transduced with different amount of the resuspend viral preparation in full medium. After 24 hrs medium was removed and replaced by fresh medium, and cells were then maintained in culture for downstream analyses.

### In vitro ubiquitination

C-terminal 3×Flag-tagged AMBRA1 and p21 were in vitro translated (IVT) using the TNT T7 Quick Coupled Transcription/Translation System (Promega) according to the manufacturer’s recommendations. In vitro ubiquitylation reaction was carried out in buffer containing 50 mM HEPES, pH 8.0, 50 mM NaCl, 1 mM DTT, 10 mM mgATP, 100 nM E1 UBE1 (RnDsystems), 1 μM UBC H5C (RnDsystems), 2.5 μM CRL4-DDB1 (RnDsystems), 0.2 nM okadaic acid, 0.2 μM NaV, and either 100 μM HA-ubiquitin (RnDsystems). Reactions were incubated at 30 °C for the indicated times, and terminated by adding NuPAGE LDS sample buffer (Thermo Fisher Scientific) followed by incubation for 5 min at 95 °C. Samples were resolved by SDS–PAGE and immunoblotted as indicated.

### Antibodies

Primary antibodies used for immunoblot (IB), immunoprecipitation (IP) immunohistochemistry (IHC), Proximity-Ligation Assay (PLA) and immunofluorescence (IF) were: α-TUBULIN (GeneTex GTX628802 - IB 1:10000), β-actin (Sigma A2066 - IB 1:5000), Myc (Santa Cruz sc-40 – IB 1:1000, IP 500 ng), HSP90 (Santa Cruz sc13119 IB -1:10000), AMBRA1 (Millipore ABC131 - IB 1:1000), AMBRA1 (G-6) (Santa Cruz sc-398204 - IB 1:1000), AMBRA1 (Novus 26190002 - 1:100), cyclin D1 (Cell Signaling 2978 - IB 1:1000), cyclin D1 (Abcam 16663 - IF 1:300 / IP 1:100), cyclin D2 (Cell Signaling - 3741 IB 1:1000), cyclin D2 (Santa Cruz sc-452 - IF 1:50), cyclin D3 (Thermo Fisher Scientific - MA5-12717 WB 1:1000), cyclin E2 (Cell Signaling 4132 - IB 1:1000), p21^Waf1/Cip1^ (Cell Signaling 2947 – IB 1:1000 / IF 1:100 / IHC 1:100, IP 250 ng), p27^Kip^^1^ (Cell Signaling 3686 – IB 1:1000, IP 250 ng), p27^Kip^^1^ (Santa Cruz sc-528 – 1:100), Ki67 (Abcam 15580 IF 1:200), HA (Y11) (Santa Cruz sc-805 - 1-500), CUL1-NEDD8 (Cell Signaling 4995 - IB 1:1000), PARP / cleaved-PARP (Abcam ab32138 - IB 1:1000), pRb 780 (Cell Signaling 8307 - IB 1:1000) pRb 807/811 (Cell Signaling 9308 - IB 1:1000), Cdt2 (Abcam ab72264 IB - 1:500), FEN1 (Novus NB100-150 PLA - 1:100), Rb (Cell Signaling 9313 IB - 1:1000), CDK4 (Cell Signaling 12790 – IB 1:1000), CDK4 (Santa Cruz sc-23896 – PLA 1:50), CDK2 (Cell Signaling 2546 – IB 1:1000), CDK2 (Santa Cruz sc-6248 – PLA 1:50), DDB1 (Novus NBP1-33061 – IB 1:1000), DDB1 (BD Biosciences 612488 – PLA 1:100), PCNA (Santa Cruz sc-56 – IB 1:1000), PCNA (Abcam ab18197 PLA – 1:100) Chk1 (Santa Cruz sc-8408 - IB 1:1000), Chk1(Ser345) (Cell Signaling 2341 - IB 1:1000), H2AX p-S139 (Merck Millipore 05-636 - IF 1:500), H2AX p-S139 (Abcam ab22551 - IB 1:500), CDK2 pT160 (Cell Signaling 2561 - IB 1:500), ATM (Abcam ab2618 – IB 1:1000), ATM pS1981 (Abcam ab81292 – IB 1:5000), p53 (Santa Cruz sc-126 – IB 1:500), UBIQUITIN (Santa Cruz sc-8017 - IB 1:500), SKP2 (Santa Cruz sc-7164 – IB 1:1000).

### Real time PCR (qRT-PCR)

RNA from cell lines was isolated by using NucleoSpin® RNA (MACHERY-NAGEL). First-strand cDNA was generated by using the GoScript Reverse Transcription System (Promega,Real-time PCR was performed using the iTAQ universal SYBR Green Supermix (Bio-Rad) or SYBR-Green (Applied Byosystems) on ViiA 7 Real-Time PCR System (Applied Biosystems) and QuantStudio 12K Flex (Applied Biosystems).

All reactions were run as triplicates. Resulting data were analyzed by t ViiA™ 7 Software. The fold changes in mRNA levels were determined relative to a control after normalizing to an internal standard as reported. The primers used in real-time PCR are the following:

*ACTIN* FW-CTGGGTATGGAATCCTGTGG / RV-GTACTTGCGCTCAGGAGGAG, *CCND1* FW-CTGGCCATGAACTACCTGGA / RV-CTCCGCCTCTGGCATTTTGG, *CCND2* FW-CACCGACTTTAAGTTTGCCA / RV-TTGGTGATCTTAGCCAGCAG

### Immunoblot analysis (IB)

Cell lysates were prepared with RIPA buffer or whole-cell lysis buffer (50 mM Tris-HCl pH 6.8, 10 % Glycerol, 2% SDS) for 5 min at 95°C at 1000 rpm agitation. All lysis buffers were added with 1x protease and phosphatase inhibitors (Sigma-Aldrich). Protein extracts were quantified using the DC protein assay (Bio-Rad), and denatured in NuPAGE® LDS Sample Buffer (Life technologies). Then, protein extracts were subjected to SDS-PAGE and transferred to polyvinylidene fluoride (PVDF) membranes in 25 mM Tris, 192 mM glycine wet electroblotting Trans-Blot turbo system (Bio-Rad laboratories), or iBlot2 Dry Blotting system (Thermo Fisher). The membranes were blocked using 5% (w/v) dry milk in PBS-Tween-20 (0.5% vol/vol) and probed with the indicated primary antibodies in blocking solution overnight at 4°C, followed by incubation in secondary horseradish-peroxidase (HRP)-conjugated antibodies (Bio-Rad) (1:5000) for 1 hour at RT. Secondary antibody detection was performed using Amersham ECL Prime (GE Healthcare) or Immobilion ECL Ultra Western HRP Substrate (Merck Millipore IBULS0100) and the signal was acquired by Invitrogen iBright CL1500 or by MP900E (Colenta) developer. Densitometric levels were quantified using Image J.

### Co-Immunoprecipitation (Co-IP)

Cells were lysed in a buffer composed of 150 mM NaCl, 0,3%CHAPS, 40mM pH 7.5 Hepes, 2 mM EDTA. Lysates (0.5 –1 mg) were then incubated at 4 °C for 30 min. Samples were precleared from unspecific bounding with 10 μl of magnetic Dyna Beads Protein G for 1 hr at 4°C. Depending on experimental conditions, equal amounts of protein were incubated with 250 ng of monoclonal anti-p21^Waf1/Cip1^ (Cell Signaling 2947) or anti-p27^Kip1^ (Cell Signaling 3686) or 500 ng anti-myc tag (Santa Cruz sc-40) overnight at 4°C, followed by 60 min incubation with 10 μl of magnetic Dyna Beads Protein G (Invitrogen).

To detect ubiquitination levels of p21^Waf1/Cip1^ and p27^Kip1^, HeLa cells overexpressing HA-tagged ubiquitin were lysed in 150 mM NaCl, 0.4 % NP-40, 10% Glycerol, 1 mM EDTA, 1 mM EGTA, 3mM MgCl2, 50 mM pH 7.4 Hepes. Lysates were then incubated at 4° for 60 min and then was added 1% SDS (Sodium-Dodecyl-Sulfate) to the final volume recovered followed by 5 min incubation at 90° C and 300 RPM. The samples were then diluted 7-fold using initial lysis buffer and equal amounts of protein were incubated with anti-p21^Waf1/Cip1^ or anti-p27^Kip1^ overnight at 4°C, followed by 60 min incubation with 10 μl of magnetic Dyna Beads Protein G (Invitrogen).

For mass-spectrometry analysis Hela wt and KO for AMBRA1 were overexpressed with either Flag p21 WT (Addgene #16240) or pMSCV-FLAG-2xSTREP-CDKN1B-Puro (Addgene #172611). For cell lysis buffer solution composed of 1% NP40, 50 mM pH 7.5 Hepes, 150 mM NaCl, protease inhibitor (Sigma) and phosphatase inhibitors NaF 5mM, Na3VO4 5mM, Beta-glycerophosphate 5mM. Lysates were then incubated at 4° C for 60 min. Flagged recombinant peptides were immunoprecipitated with Pierce™Anti-DYKDDDDK Magnetic Agarose (Product No. A36797) and then eluted with Pierce™ 3x DYKDDDDK Peptide (Product No. 36805), according to manufacturer instructions.

### LC-MS/MS Analysis

Instruments for LC-MS/MS analysis consisted of a NanoLC 1200 coupled via a nano-electrospray ionization source to the quadrupole-based Q Exactive HF benchtop mass spectrometer (Michalski et al., 2011). For the chromatographic separation, a binary buffer system consisting of solution A: 0.1% formic acid and B: 80% acetonitrile, 0.1% formic acid was used. The peptides were separated according to their hydrophobicity on an analytical column (75 μm) in-house packed with C18-AQ 1.9 μm C18 resin with a gradient of 7–32% solvent B in 45 min, 32–45% B in 5 min, 45–95% B in 3 min, 95–5% B in 5 min at a flow rate of 300 nl/min. MS data acquisition was performed in DIA (Data Independent Acquisition) mode using 32 variable windows covering a mass range of 300–1650 m/z. The resolution was set to 60’000 for MS1 and 30’000 for MS2. The AGC was 3e6 in both MS1 and MS2, with a maximum injection time of 60 ms in MS1 and 54 ms in MS2. NCE were set to 25%, 27.5%, 30%.

All acquired raw files were processed using Spectronaut 18 software. For protein assignment, spectra were correlated with the Human data base (v. 2023). Searches were performed with tryptic specifications and default settings for mass tolerances for both MS and MS/MS spectra. The other parameters were set as follow:

- Fixed modifications: Carbamidomethyl (C)
- Variable modifications: Oxidation, Acetyl (N-term)
- Digestion: Trypsin, Lys-C
- Min. peptide length = 7Da
- Max. peptide mass = 470Da
- False discovery rate for proteins and peptide-spectrum = 1%

The Perseus software (1.6.2.3) was used to logarithmize, group and filter [after filter] the protein abundance. ANOVA and t-test analysis were performed setting FDR = 0.05. Proteins with difference. ANOVA q-value or Log2 Difference≥ ± 0.5 and q-value <0.05 were considered significantly enriched (see “Significantly deregulated” column in the excel file).

### Flow cytometry and FACS Sorting

Stable cell lines expressing the FUCCI CA (5) plasmid reporter were washed with PBS-1X, trypsinized and collected in DMEM without phenol red (Gibco, Thermo-Fisher Scientific). Cells were filtered using a 40 µm Cell Strainer (Falcon) and kept on ice. Cell cycle analysis and sorting of cell cycle phases subpopulations were carried out by the FACS Celesta (BD bioscience) using the FITC filter to detect cells in S phase, RFP+ for cells in G1 and cells showing double-positive signal (FITC+/RFP+) were considered in G2/M phase (Supplementary Information). For cell sorting a FACS Aria cell sorter was used and at least 1×10^5 cells for each cell cycle phase were sorted. Cells were collected by centrifugation and the pellet was utilized for Western blotting samples preparation.

### PLA - Proximity Ligation Assay

DAOY cells were plated in fibronectin coated 13mm slides. Cells were then transiently interfered for AMBRA1 expression for 48 hrs before being fixed with 4 % PFA. Coverslips were then processed with Duolink® Proximity Ligation Assays (PLA) Kit (Sigma-Aldrich) according to manufacturer instructions and lastly stained using Duolink In Situ Mounting Media with DAPI for 15 mins. Samples were mounted on the slide and images were acquired with Olympus Confocal Microscope and CellDiscoverer7 at 20x and foci detection was performed through a customized pipeline in Arivis Vision4d software 4.1.1.

### Annexin V staining

Cells were washed twice with cold PBS and then were resuspended in 1X Annexin V Binding Buffer (BD Pharmigen #51-55121E). 1 x 10^5^ cells were transferred into 5 ml round bottom tubes. 5 µl of APC Annexin V (BD Pharmigen #550475) were added to each culture tube and cells were incubated for 15 min at RT in the dark. The tubes were analyzed by flow cytometry immediately after.

### Cell viability assay

To obtain the total cell count, cells were stained with CellTracker 488 (Invitrogen) for 30 min at 37 °C and analysed using a Celigo imaging cytometer (Nexcelom).

### UV irradiation and pharmacological treatments

HeLa wt and KO for AMBRA1, DAOY and ONS76 cells were exposed to UV radiation of either 20, 40 or 60 J/m^2^, according to experimental conditions using an UV Stratalinker 2400. For protein stability evaluation cells were treated with 100 μg/mL CHX (cycloheximide - SIGMA C4859) alone or in combination with 5 μM MG132 (SIGMA M7449) for the indicated time points. NEDD-8 activating enzyme inhibitor, MLN4924, for Cullins inhibition was used in DAOY, ONS76, HeLa wt and KO for AMBRA1 for 4 hrs with 2,5 μM. Cells were treated with AZD7762 (SML0350 Sigma) 200 nM, FEN1i 10 µM. Medulloblastoma cell lines were treated with either abemaciclib or palbociclib for 24 hrs at the indicated doses. HCT-116 2×Flag-mAID-AMBRA1 cell line was treated with Doxycyclin 1µg/ml (D9891, Sigma), Auxin 0,5 mM (I5148, Sigma) and MS28 6 µM (Xiong, Y. et al., 2022) for the time points indicated in figure legends.

### Quantitative image-based cytometry (QIBC)

For immunocytochemistry, cells were grown on 96-well (Greiner Screenstar 655866) 24h before the transfection. 48 hrs after the initial silencing cells were incubated with 10 µM EdU for 30 min at 37 °C before were fixed in 4% paraformaldehyde in PBS for 15 min and washed 3 times in PBS. Permeabilization was performed in PBS plus 0.5 % Triton X-100 for 5 min at room temperature (RT), followed by 30 min blocking in the blocking buffer (PBS plus 5 % FBS, 1 % BSA and 0.2 % Triton X-100). Click-iT EdU Imaging Kits (Invitrogen) were used for EdU detection according to the manufacturer’s instructions. Cells were incubated with primary antibodies in a blocking buffer for 2 hrs at RT and washed 3 times in PBS plus 0.1 % Triton X-100. Then, cells were incubated with the appropriate secondary antibodies conjugated to Alexa Fluor 488 or Alexa Fluor 568 (Life Technologies) diluted in a blocking buffer for 45 mins at RT. DNA was stained using Hoechst 33342 (ThermoFisher Scientific, H3570, 1:1000 in PBS) 10 min at RT. Automated multichannel wide-field microscopy for high-content imaging and quantitative image-based cytometry was performed using CellDiscoverer7 (Zeiss) using 20x magnification objective. After background subtraction, cells were segmented for the cytoplasmic and nuclear areas with Arivis software using a customized pipeline implemented with cellpose algorithm (Stringer et al., 2021). For CDK2 and CDK4/6 activity analysis cells were segmented using the same pipeline. CDK4/6 activity correction based on the activity of the CDK2 reporter was performed as described previously (Yang et al., 2020). For each condition, 6×6 tiles of 3 independent experiments were acquired and at least 1,000 cells per replicate were processed using Zen 2.6 (Blue Edition) software (Zeiss). Scatter plots and bubble plots were generated with the ggplot2 package in R.

### DNA fibers analysis

Cell cultures transfected with siRNA and/or treated with different drugs were pulse-labeled with 25 μM of CldU (Sigma-Aldrich) for 20 min, followed by a gentle wash with fresh pre-warmed media and the second pulse of 250 μM of IdU (Sigma-Aldrich) for 20 min. Labeled cells were harvested and DNA fiber spreads prepared as previously described^42^. For every single experimental condition, 5 slides were stretched and 2-3 slides for each condition were stained. CldU was detected first with the rat anti-BrdU antibody (Serotec, OBT0030) and IdU with the mouse anti-BrdU antibody (Becton Dickinson, 347580). Secondary antibodies were DyLight 550 anti-rat (Thermo Fisher Scientific) and Alexa Fluor 488 anti-mouse (Invitrogen), respectively. Images of well-spread DNA fibers were acquired using a LSM800 confocal microscope (Carl Zeiss) and a Plan-Apochromat 63x/1,4 N.A. oil immersion objective (Carl Zeiss). Images were acquired semi-automatically by using software autofocus and tile-arrays. Double-labeled replication forks were analyzed manually using LSM ZEN software. Between 100 and 250 replication forks were scored for each slide, and fork measurements for all slides for the same experimental condition were pooled together. At least one additional independent experiment was performed, and, if the experiments did not differ statistically, the total number of DNA fibres from all experiments is presented.

### S1 nuclease assay for DNA fibers

Cell cultures in 6 cm dishes transfected with siRNA were pulse-labeled with 250 μM of ldU (Sigma-Aldrich) for 60 min, followed by a gentle wash with PBS. Cytoskeletal disruption was achieved using 0.5 ml of fresh CSK100 (100mM NaCl, 10mM MOPS, 3mM MgCl2, 300mM sucrose, 0.5% Triton X-100) for 10 min at RT. After a careful wash with PBS and equilibration using the S1 nuclease buffer (30mM sodium acetate, 10mM zinc acetate, 5% glycerol, 50mM NaCl, pH 4.6), the cells were treated with 20 U/ml S1 nuclease for 30 min at 37°C. After S1 nuclease removal, nuclei were harvested using a cell scraper and 0.5 ml of 0.1% BSA in PBS. Nuclei were kept on ice for subsequent DNA fiber spreads (adapted protocol from Quinet et al. 2017).

### Live Imaging

For U2OS-FUCCI time-lapse imaging, cells were plated at low density (approximately 10,000 cells per well) and imaged in 96-well plates in DMEM inside a heated 37°C chamber with 5% CO2. Images were taken every 30 min with a 20×/0.5 NA air objective using a Zeiss Cell Discoverer 7. For cell tracking ImageJ plugin TrackMate^57^ with StarDist segmetation was used.

### Histology and Immunohistochemistry

Immunofluorescence (IF) analysis of tissue sections of the Ambra1/Nestin-Cre mouse model were carried out on cryoembedded embryos (OCT). Before the embedding embryos and pups brain were fixed 24h in 4% formaldehyde in PBS, then cryoprotected with 30% sucrose in PBS solution for 48h. After sinking, tissues were enclosed in 1:1 mixture of OCT (Thermo Fischer) and 30% sucrose/PBS solution, embedded in cryomolds that were pre-chilled in an isopentane bath on dry ice and stored at -80°C. Lastly, tissues were cut with Leica CM3050S cryostat and placed on SuperFrost Plus glasses (Thermo Fischer). For IF, sections were washed and rehydrated with PBS, permeabilized and blocked with 0.3% Triton-X in Protein Block (Abcam) solution. The slides were incubated with primary antibodies in Protein Block overnight at 4°C in a humidified chamber. Sections were then washed three times in PBS and incubated with the appropriate secondary antibodies conjugated to Alexa Fluor 488 or Alexa Fluor 568 (Life Technologies) diluted in Protein Block for 2 hrs at RT. Eventually, sections were stained with Hoechst 33342 for 10 min at RT. The slides were mounted with Fluoromount (Sigma F4680). IF images were taken with Hamamatsu Nanozoomer S60 or Confocal Microscope Olympus FV1000. For H/E of MB-SHH patients, sections were then dipped in Gill’s Hematoxylin No. 2 Solution (Bio-Optica, Cat. No. 05-06014/L) for 30 seconds. Sections were washed in ddH2O, followed by 0.5% alcoholic eosin (Diapath, Cat. No. C0353). Sections were dehydrated with one, 10-second wash in 90% ethanol, and three, 10-second washes with 100% ethanol. Lastly, sections were immersed in Diasolv (Diapath, Cat. No. H0315) for three times.

All samples of human tissue research in this study were collected at the Bambino Gesù Children’s Hospital in Rome, with Institutional Review Board approval. These clinical MB specimens were examined and diagnosed by pathologists. Tissue sections of clinical specimens were stained with antibodies against AMBRA1, cyclin D1 and p21^Waf1/Kip1^. The analyses were assessed by an experienced pathologist (S.R.), blinded to sample. Digital images were acquired by Hamamatsu NDP Nanozoomer digital pathology slidescanner.

### Analyses of publicly available transcriptomics data

We used the R2 genomics platform (http://r2.amc.nl) to identify suitable transcriptomics datasets with molecular and histological subtypes available for MB. We selected the following datasets: Pomeroy (204 samples, MAS5.0 normalized, u133a chip) and Cavalli (763 samples, rma_sketch normalized, hugene11t chip). For each of them, we download log2 transformed and normalized data through the data grabber function for each subgroup (normal cerebellum, SHH) and subtype (SHH - α, β, γ, δ) for AMBRA1 (52731_at probe) expression levels. We employed a parametric Welch’s T-test to assess the significance of the differences in expression levels of the same gene in the different subgroups and subtypes.

### DepMap co-occurrency analysis

Public 22Q2 CERES corrected gene effects of CRISPR KO were downloaded from the Depmap portal (https://depmap.org/portal/). Correlation analysis was performed and Top500 genes were isolated for AMBRA1 co-dependencies. Following pathway analysis performed through ConsensusPathDB-human and top 15 Reactome pathways were plotted.

### Kaplan-Meier survival

Survival analysis was generated using the dataset GSE85217 (Cavalli). AMBRA1 gene expression levels were expressed as a binary variable using the cut-off value generated by the Kaplan Scan online algorithm (https://hgserver1.amc.nl/cgi-bin/r2/main.cgi?option=kaplanscan1). The overall survival right-censored at 5 years, was estimated using the Kaplan–Meier method and p-values were assessed using the log-rank test to express statistical differences in overall survival among patients in MB-SHH).

## Data availability

Kaplan–Meier analysis in Fig. 1b and referenced during the study are available in a public repository from the websites (https://r2.amc.nl/). Co-dependencies analysis in Fig. 1a, 6b and 6f, are available in the public repository DepMap (depmap.org)

Representative gating strategy of FACS analysis are included in **Supplementary Fig. 1**.

## Statistical analysis and Data reproducibility

Statistical analyses were performed with Prism software (GraphPad Software) or with R environment. All statistical parameters including the exact value of *n*, type of replicates, the statistical test, error bars and significance are reported in all associated figure legends. All experimental findings were verified in ≥3 independent experiments.

No statistical methods were used to pre-determine the sample size. The experiments were not randomized, and the investigators were not blinded to allocation during experiments and outcome assessment.

All the data behind the statistical analysis will be provided as **Source Data**.

---

**Extended Data Figure 1.**
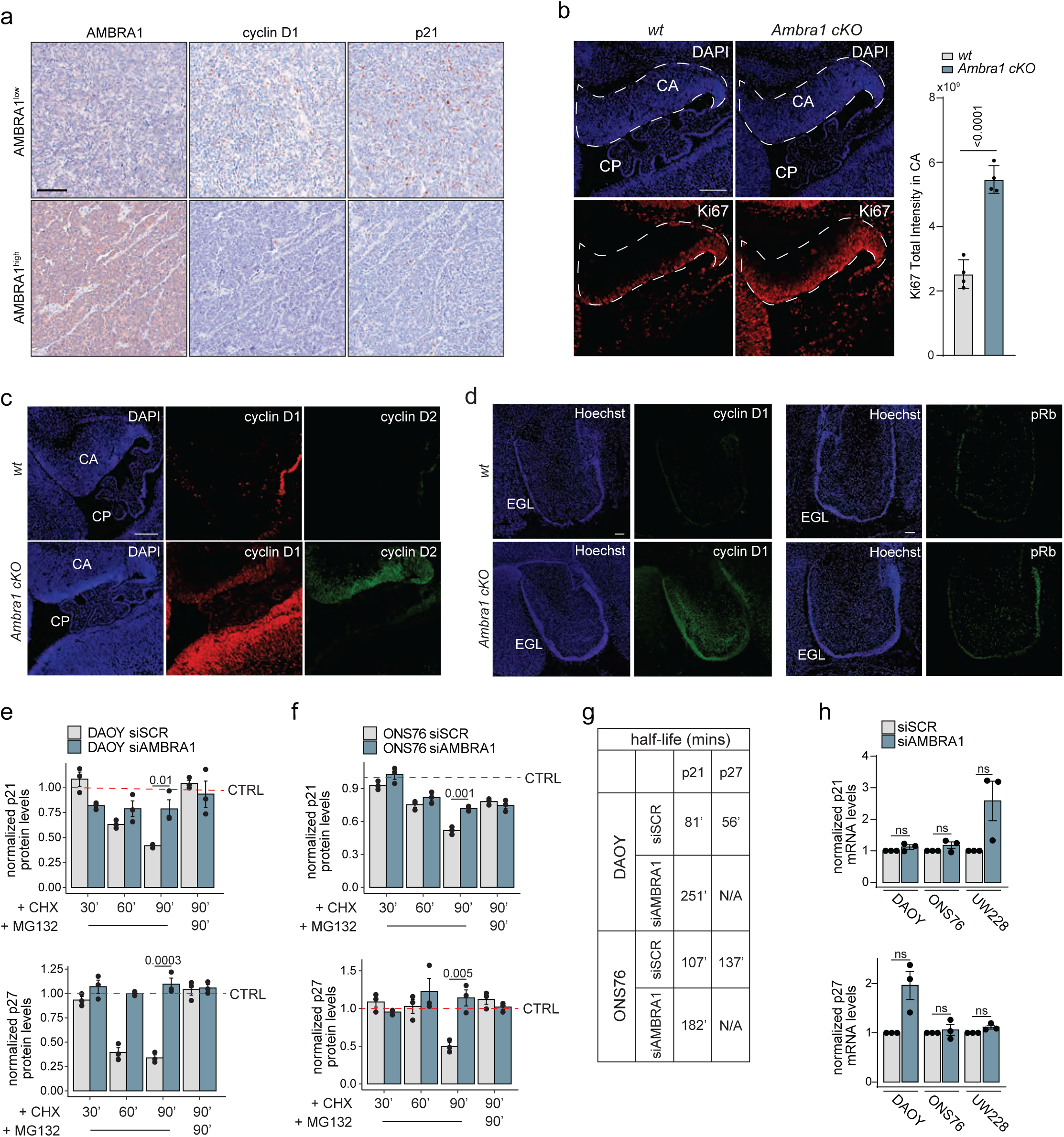
Ambra1 controls cerebellar development through cell cycle regulation. **a**, Representative images of IHC for the indicated proteins in MB-SHH patients showed in Fig. 4d (n=14). Scale bar: 100 um. **b**, Sagittal sections from wild-type and Ambra1 cKO E13.5 embryos (*Top panel*), stained for Ki67 and Hoechst (n=4). Dashed area refers to CA, cerebellar anlage; CP, choroid plexus. *Right panel*, Barplot of Ki67 total intensity in the CA. **c,** Sagittal sections from wild-type and Ambra1 cKO E13.5 embryos stained for cyclin D1, cyclin D2 and Hoechst (n=4). **d**, Sagittal sections from wild-type and Ambra1 cKO E18.5 embryos stained for cyclin D1 (*Left panel*), pRb-S807/811 (*Right panel*) and Hoechst (n=3 – pRb; n=4 – cyclin D1). **e-f**, Barplot representing over-CTRL normalized quantification of p21 (*Top panel*) and p27 (*Bottom panel*) protein levels in DAOY (**e**) and ONS76 (**f**) cells in control or AMBRA1-depleted conditions (shown in Fig 1i**-j**), and presented as mean value ±SD. **g**, Schematic table of half-life measurements calculated using exponential decay for the indicated proteins in control and AMBRA1-depleted MB cells.. **h**, Barplot of quantitative PCR with reverse transcription (qRT‒PCR) of the indicated genes in the three MB-SHH cell lines. Unless otherwise stated data are presented as mean value ±SEM and n refers to biological independent samples. Data were analyzed using unpaired Student t-test (**b**, **e-f**) and Wilcoxon Test (**g**).

**Extended Data Figure 2.**
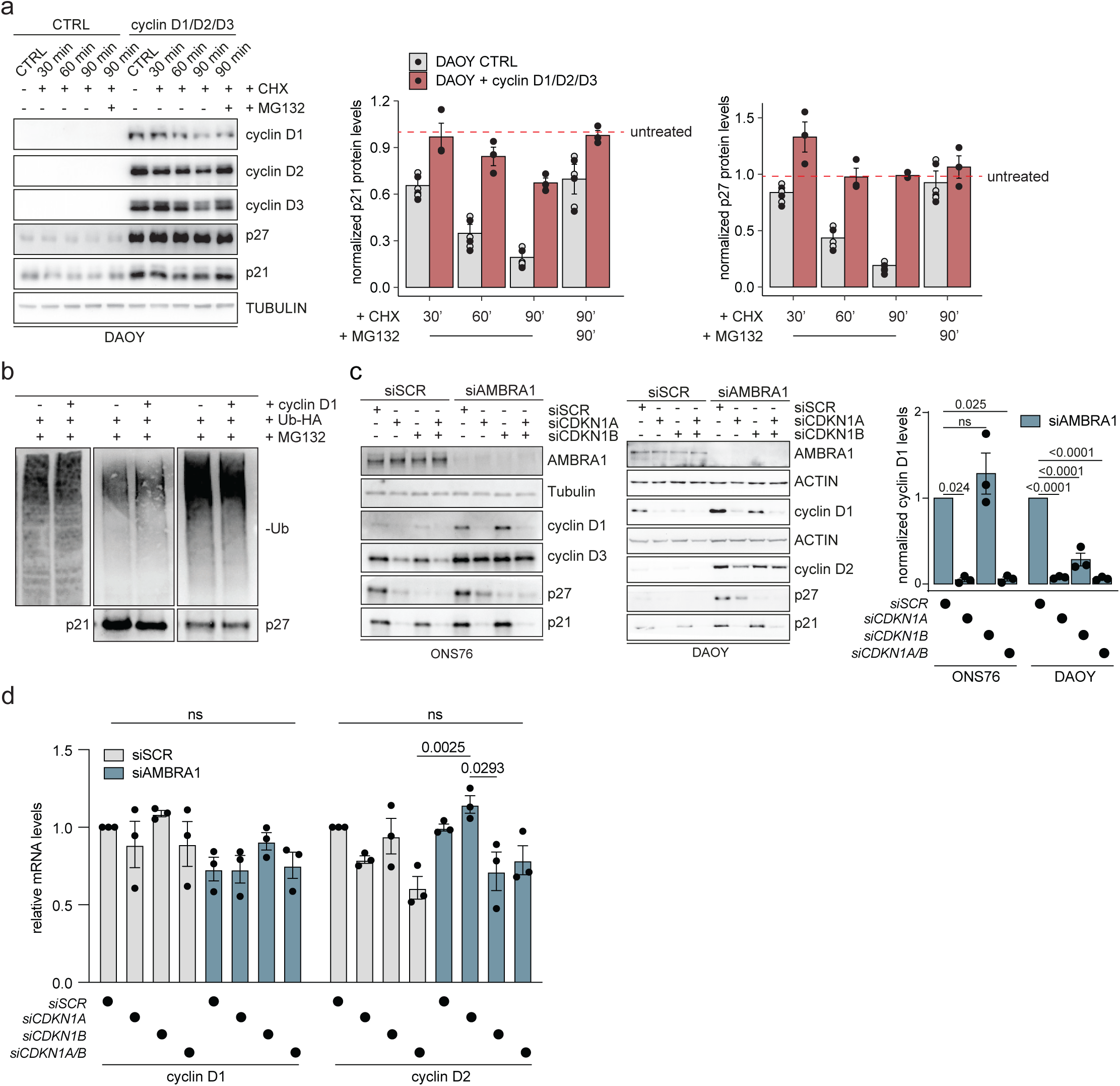
p21 levels affect cyclins D stability. **a**, (*Left panel*) IB of the indicated proteins in DAOY cells overexpressing the three D-type cyclins, treated with 100 µg/ml cycloheximide (CHX) and/or 10 µM MG132 for the indicated time points. Corresponding barplots representing over-untreated normalized quantification of p21 (*Middle panel*) and p27 (*Right panel*) protein levels (n=3). Data are presented as mean value ±SD **b**, IB for the *in vivo* ubiquitination levels of p21 and p27. Hela cells constitutively overexpressing a doxycycline-inducible form of cyclin D1 were transfected to overexpress a plasmid encoding for an HA-tagged form of ubiquitin and treated with 5 µM MG132 before harvesting as indicated (n=3). **c**, IB for the indicated proteins in ONS76 (*Left panel*) and DAOY (*Middle panel*) cells silenced or not for the indicated genes (n=3). (*Right panel*) Corresponding barplot of *siSCR*-normalized protein levels of cyclin D1 in AMBRA1-silenced cells. **d**, Barplot of normalized mRNA levels of *CCND1* and *CCND2* by q-PCR in DAOY treated as in **c** (n=3). Unless otherwise stated data are presented as mean value ±SEM and n refers to biological independent samples. Data were analyzed using unpaired Student t-test (**b**) or One-way ANOVA (**c-d**).

**Extended Data Figure 3.**
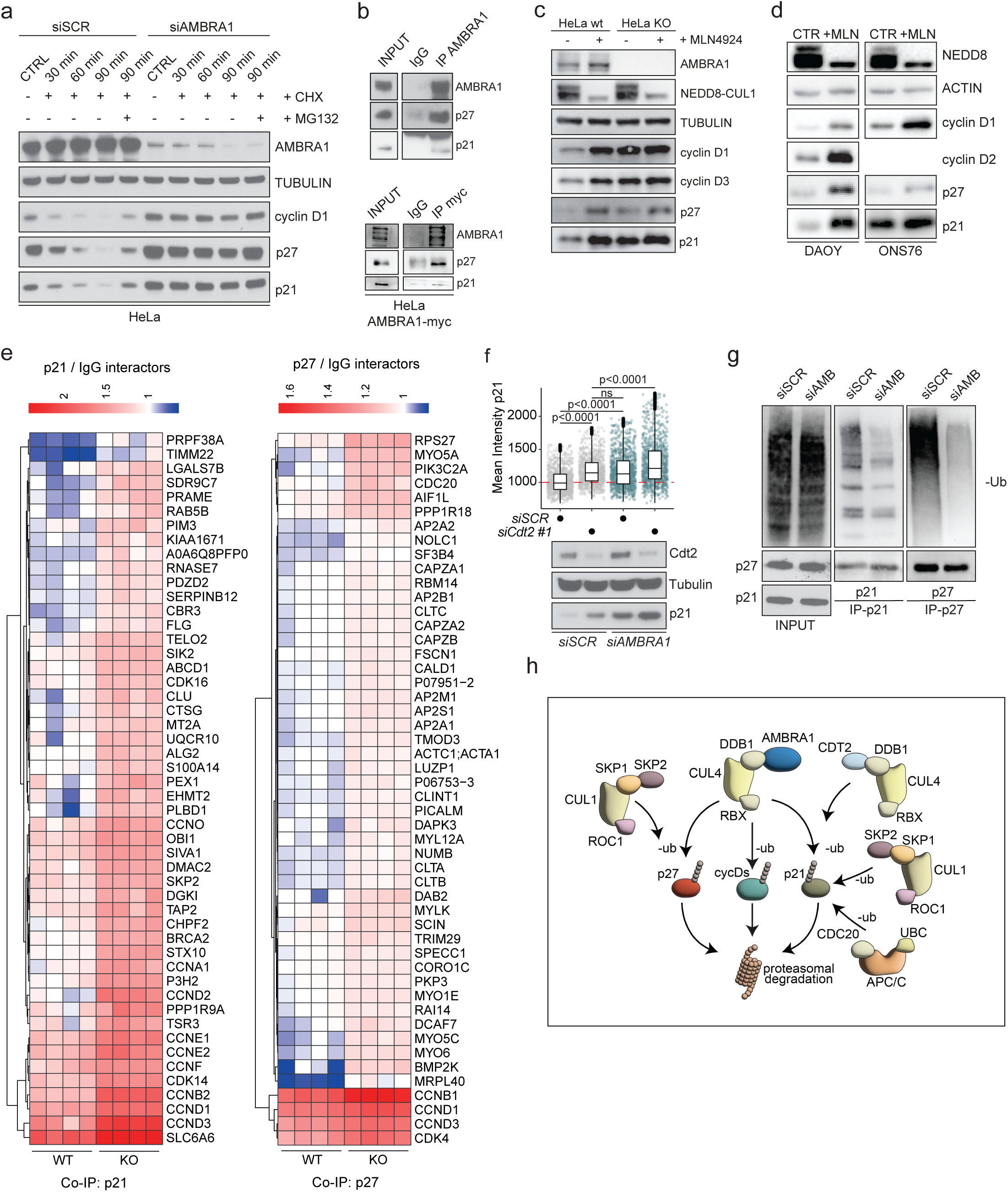
AMBRA1 depletion impairs DDB1-mediated degradation of p21 and p27. **a**, IB of the indicated proteins in HeLa cells depleted for AMBRA1 with siRNA and treated with 100 µg/ml cycloheximide (CHX) and/or 10 µM MG132 for the indicated time points (n=3). **b**, IB of the indicated proteins for the co-IP of endogenous (*Top panel*) and exogenous (*Bottom panel*) expressed AMBRA1 (n=3). **c-d**, IB of the indicated proteins in wt or *AMBRA1*-KO HeLa cells (**c**) and in DAOY and ONS76 cells silenced for scramble or AMBRA1 (**d**), treated or not with NEDD8 inhibitor 2,5 µM MLN4924 for 4 hours (hrs) (n=3). **e**, Heatmap of top50-upregulated interactions (log2FC) from proteomics analysis in wt and *AMBRA1*-KO HeLa cells overexpressed with either p21 (*Left panel*) or p27 (*Right panel*) Flag-tagged plasmids, normalized over the respective IgG control samples (n=4). **f**, QIBC (*Top panel*) and IB (*Bottom panel*) analysis for the indicated proteins in DAOY cells silenced or not for the indicated genes (QIBC: n=1000 in 3 independent technical replicates; IB: n=2). **g**, IB for the *in vivo* ubiquitination levels of p21 and p27. DAOY cells deplete or not for *AMBRA1* were transfected to overexpress a plasmid encoding for a HA-tagged form of ubiquitin and treated with 5 µM MG132 before harvesting as indicated (n=2). **h**, Graphical representation illustrating the various pathways involved in the degradation of p21, p27 and D-type cyclins ubiquitylation and degradation. Unless otherwise stated data are presented as mean value ±SD and n refers to biological independent samples. Data were analyzed using One-way ANOVA (**f**).

**Extended Data Figure 4.**
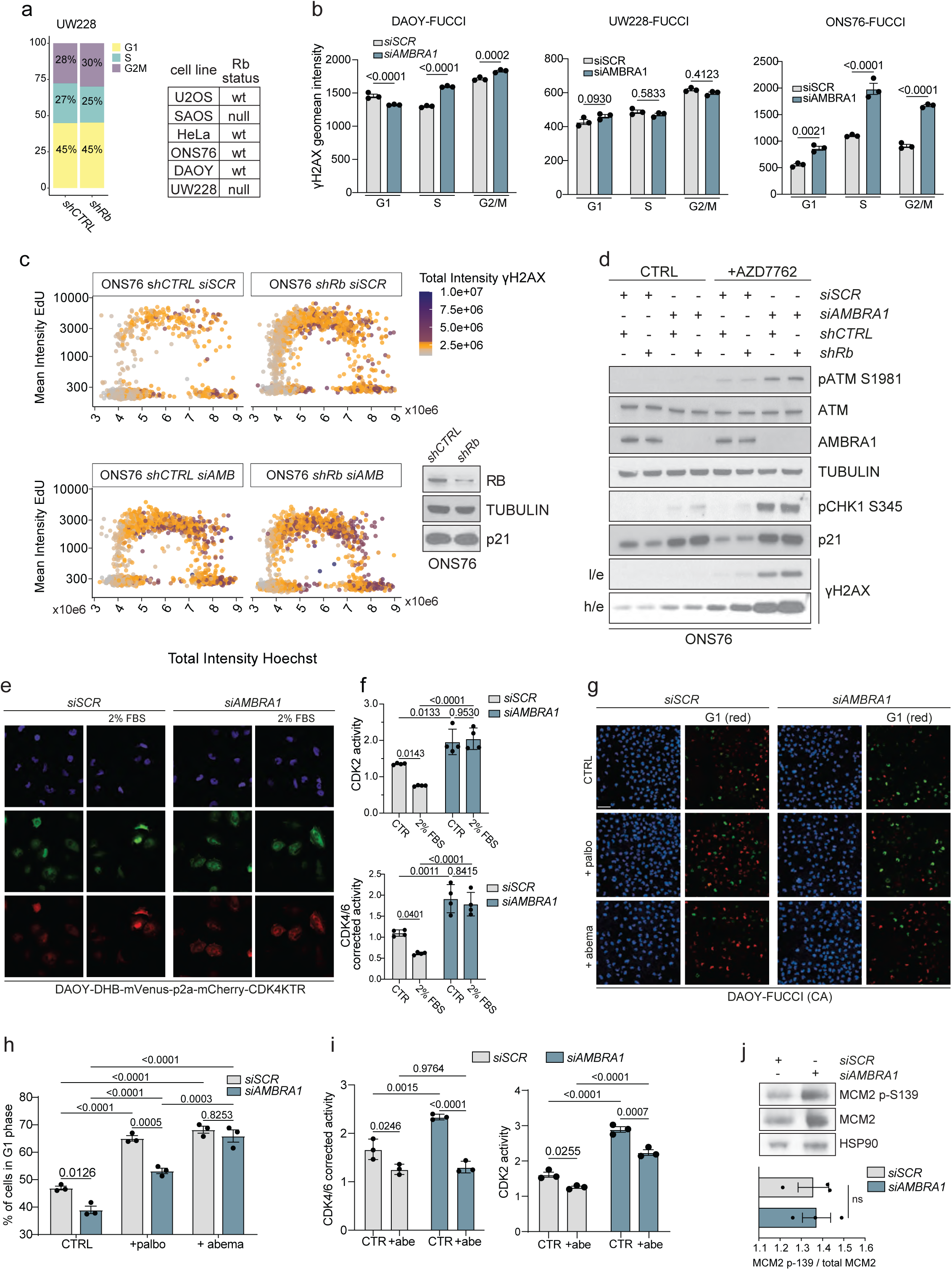
DNA damage induction following AMBRA1 depletion is partially independent of altered G1/S transition. **a,** (*Left panel*) Stacked barplot of UW228 in control condition constitutively downregulated for Rb gene expression (*shRb*) (n = 400 in 3 biological independent replicates). (*Right panel*) Table summarizing Rb status across the cell lines used in this study. **b**, Barplot of γH2AX geomean intensity analysed by flow cytometry in DAOY-FUCCI (CA), UW228-FUCCI (CA) and ONS76-FUCCI (CA) in control or AMBRA1-depleted condition (n=10000 events in three independent technical replicates). **c**, (*Left panel*) QIBC analysis of γH2AX total intensity along cell cycle of control (*shCTRL*) or *shRb* ONS76 cells in scramble or AMBRA1-depleted condition stained for EdU and Hoechst (2000 cells are displayed per condition). (*Right panel*), IB for the indicated proteins of whole-cell extracts from of control (*shCTRL*) or *shRb* ONS76. **d**, IB for the indicated proteins of whole-cell extracts from of control (*shCTRL*) or *shRb* ONS76 cells in scramble or AMBRA1-depleted condition, untreated (CTRL) or treated with 200 nM AZD7762 for 24 hrs (n=3). h/e: high exposure time; l/e: low exposure time. **e**, (*Left panel*) Representative images from DAOY cells engineered to constitutively express a CDK4/6 (red) and CDK2 (green) activity reporter. Barplot of CDK2 activity (*Top Right panel*) and CDK4/6 corrected activity (*Bottom Right panel*). Cells were depleted for scramble or AMBRA1 (siRNA) and FBS-deprived (FBS 2%) or not for 24 hrs and then analysed by high-content imaging (n=4). Scale bar 100 µm. **g**, Representative images of DAOY-FUCCI (CA) cells treated for 24 hrs with 1µM palbociclib and 500 nM abemaciclib and analysed by high-content imaging. Scale Bar: 100 µm. **h**, Barplot for G1 phase cells % analysed in **g** (n=3). **i**, Barplot for CDK4/6 and CDK2 (green) activity reporter. Cells were depleted for scramble or AMBRA1 (siRNA) and then treated or not with 500 nM abemaciclib for 24 hrs and then analysed by high-content imaging (n=3). **j**, (*Top panel*) IB for the indicated proteins of whole-cell extracts in ONS76 cells silenced or not for the indicated genes. (*Bottom panel*) Barplot of corresponding quantification of normalized phosphorylated levels over total (n=3). Unless otherwise stated data are presented as mean value ±SEM and n refers to biological independent samples. Data were analysed using One-way ANOVA (**b**, **f**).

**Extended Data Figure 5.**
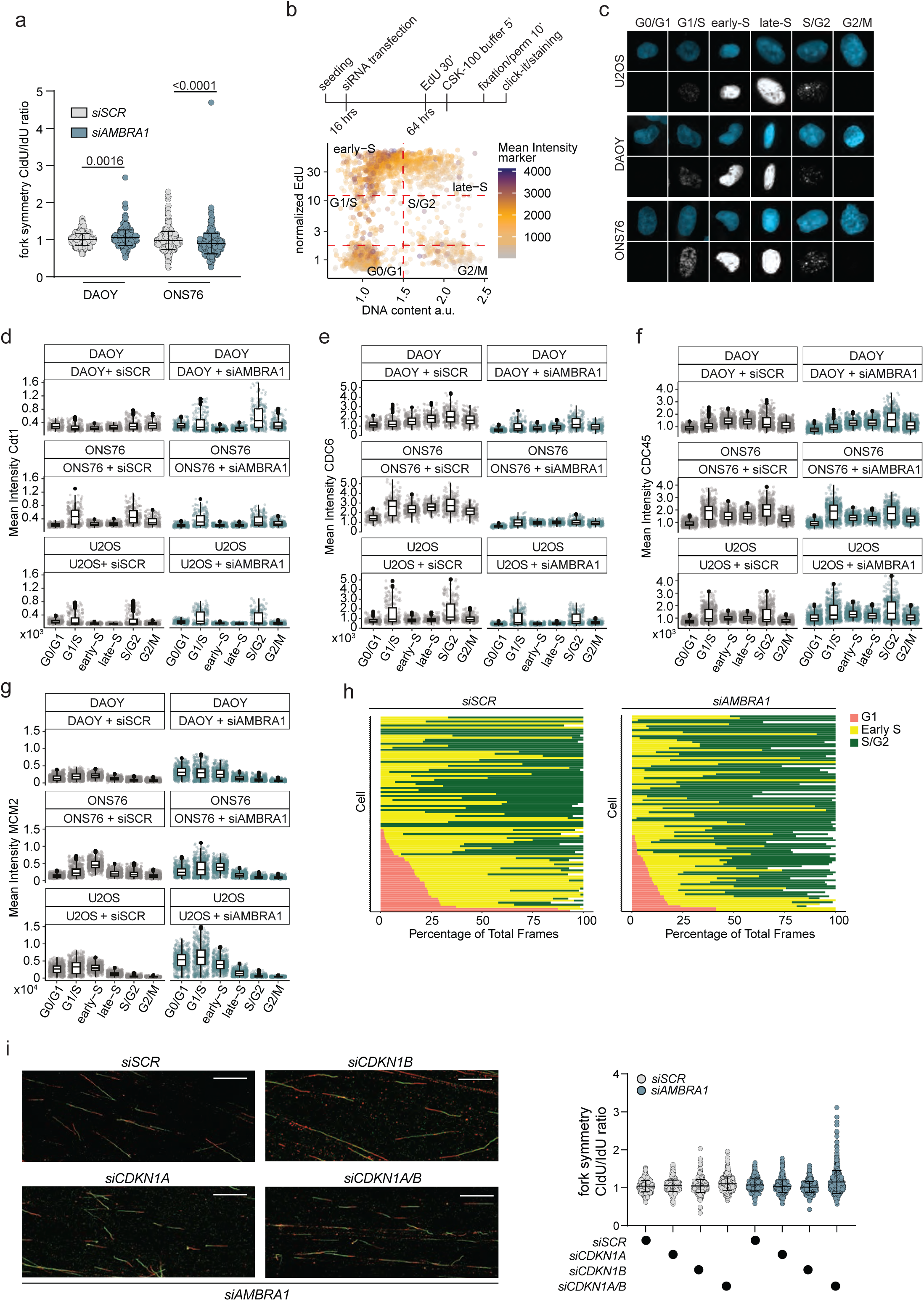
AMBRA1 depletion alters replisome processivity without affecting origin firing. **a**, Replication fork simmetry from DAOY and ONS76 cells depleted or not for AMBRA1 (DAOY *siSCR*: n = 546; DAOY *siAMBRA1*: n = 537 / ONS76 *siSCR*: n = 576; ONS76 *siAMBRA1*: n = 552). **b**, Top panel: Timeline of a representative experimental workflow utilizing Quantitative Image-Based Cytometry (QIBC) technology. Bottom panel: Following image acquisition, nuclei or cells are segmented using a customized pipeline based on the Cellpose algorithm. Segmented single-cell data are then visualized in scatter plots according to DNA content and EdU incorporation levels. **c**, Representative fluorescence images of U2OS ans MB-SHH cells nuclei stained with DAPI (light blue) and EdU (white), illustrating nuclear morphology and DNA replication activity with the corresponding cell cycle phase. **d-g**, e, QIBC analysis of chromatin-bound Cdt1 (**d**), CDC6 (**e**), CDC45 (**f**) and MCM2 (**g**) in U2OS, DAOY and ONS76 cells, stained for EdU and Hoechst (CDC6, CDC45, MCM2: 500 cells are displayed per cell cycle phase, Cdt1: 200 cells are displayed per cell cycle). **h**, Cell cycle profile of individual U2OS-FUCCI cells (each bar represents one cell) 24 hrs after AMBRA1 silencing imaged 48 hrs (n=100 randomly selected from 3 independent technical replicates). **i**, (*Left panel*) Representative images of replication forks with the corresponding analysis (*Right panel*) of replication fork speed (Fig. 5d) and symmetry (*Left panel*) in U2OS cells treated as in Fig. 4c (U2OS *siSCR* + *siSCR*: n = 513; U2OS *siSCR* + *siCDKN1A*: n = 561; U2OS *siSCR* + *siCDKN1B*: n = 516; U2OS *siSCR* + *siCDKN1A*/*B*: n = 564; U2OS *siAMBRA1* + *siSCR*: n =533; U2OS *siAMBRA1* + *siCDKN1A*: n = 527; U2OS *siAMBRA1* + *siCDKN1B*: n = 540; U2OS *siAMBRA1* + *siCDKN1A*/B: n = 564; in three technical independent experiments).

**Extended Data Figure 6.**
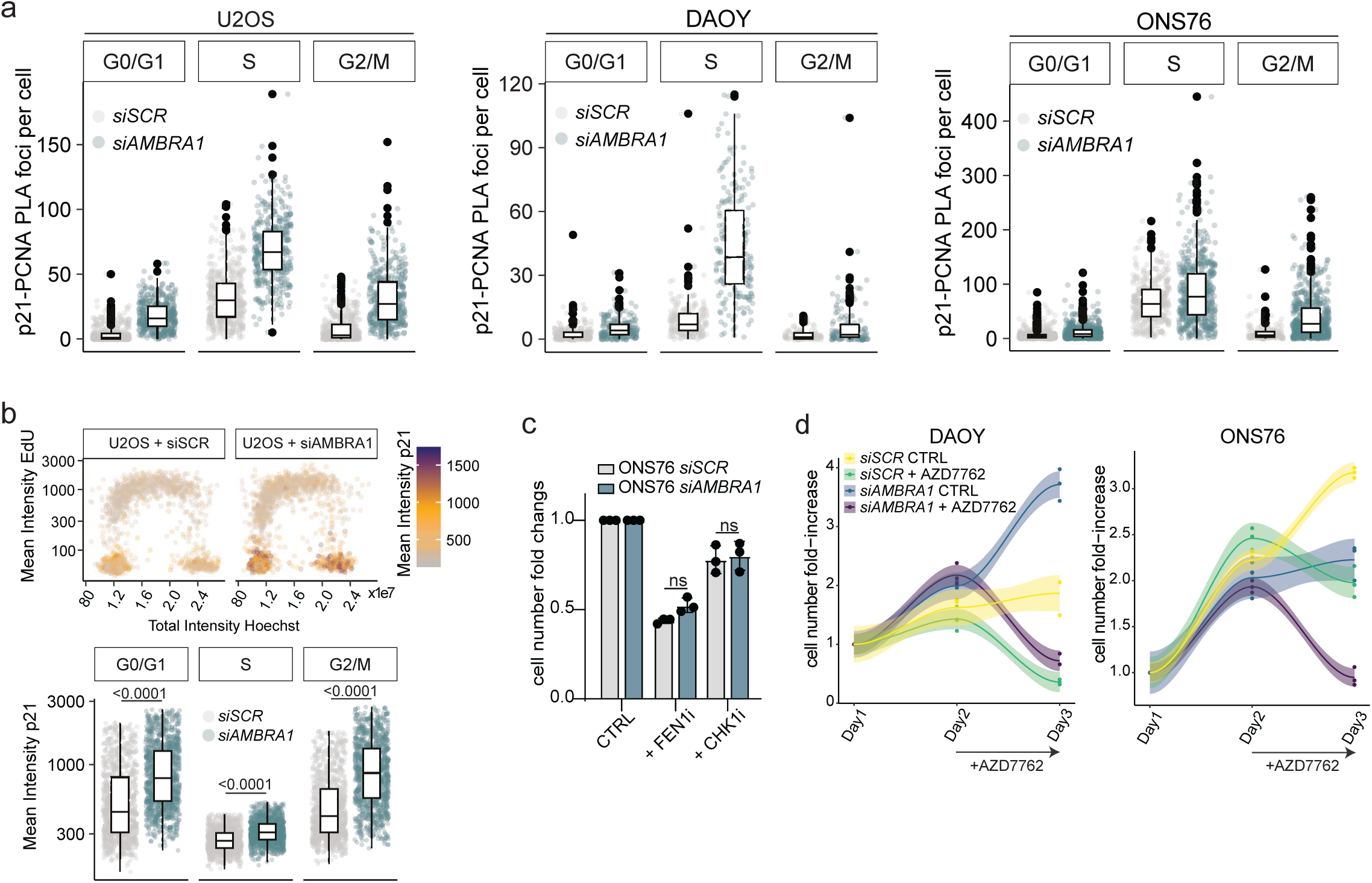
AMBRA1 depletion sensitizes to lagging strand synthesis inhibition. **a**, Boxplot (median and IQR) of QIBC analysis of PLA reaction in pre-extracted U2OS, DAOY an ONS76 cells treated as in Fig. 5e (U2OS n = 400 for all cell cycle phases; DAOY *siSCR*: G1 = 400; S = 400; G2/M = 110; DAOY *siAMBRA1*: G1 = 400; S = 400; G2/M = 118; ONS76 *siSCR*: G1 = 400; S = 400; G2/M = 227; ONS76 *siAMBRA1*: G1 = 400; S = 400; G2/M = 537 in three technical independent experiments). **b**, QIBC analysis of control and AMBRA1-depleted U2OS cells stained for p21, EdU and Hoechst. (*Top panel*) Scatter plot for cell cycle distribution and p21 nuclear intensity (2000 cells are displayed per the condition of three independent replicates). (*Bottom panel*) Boxplot (median and IQR) of p21 nuclear intensity (n = 1000 for all cell cycle phases). **c,** Barplot representing fold-change cell number in ONS76 cell line depleted or not for AMBRA1, treated for 48 hrs with the indicated cytotoxic compounds. **d**, Fold-change cell number increase in DAOY (*Left panel*) and ONS76 (*Right panel*) cell line depleted or not for AMBRA1, treated for 24 hrs with the indicated cytotoxic compound. Unless otherwise stated data are presented as mean value ±SD and n refers to biological independent samples. Data were analysed using One-way ANOVA (**b**, **c**).

**Extended Data Figure 7.**
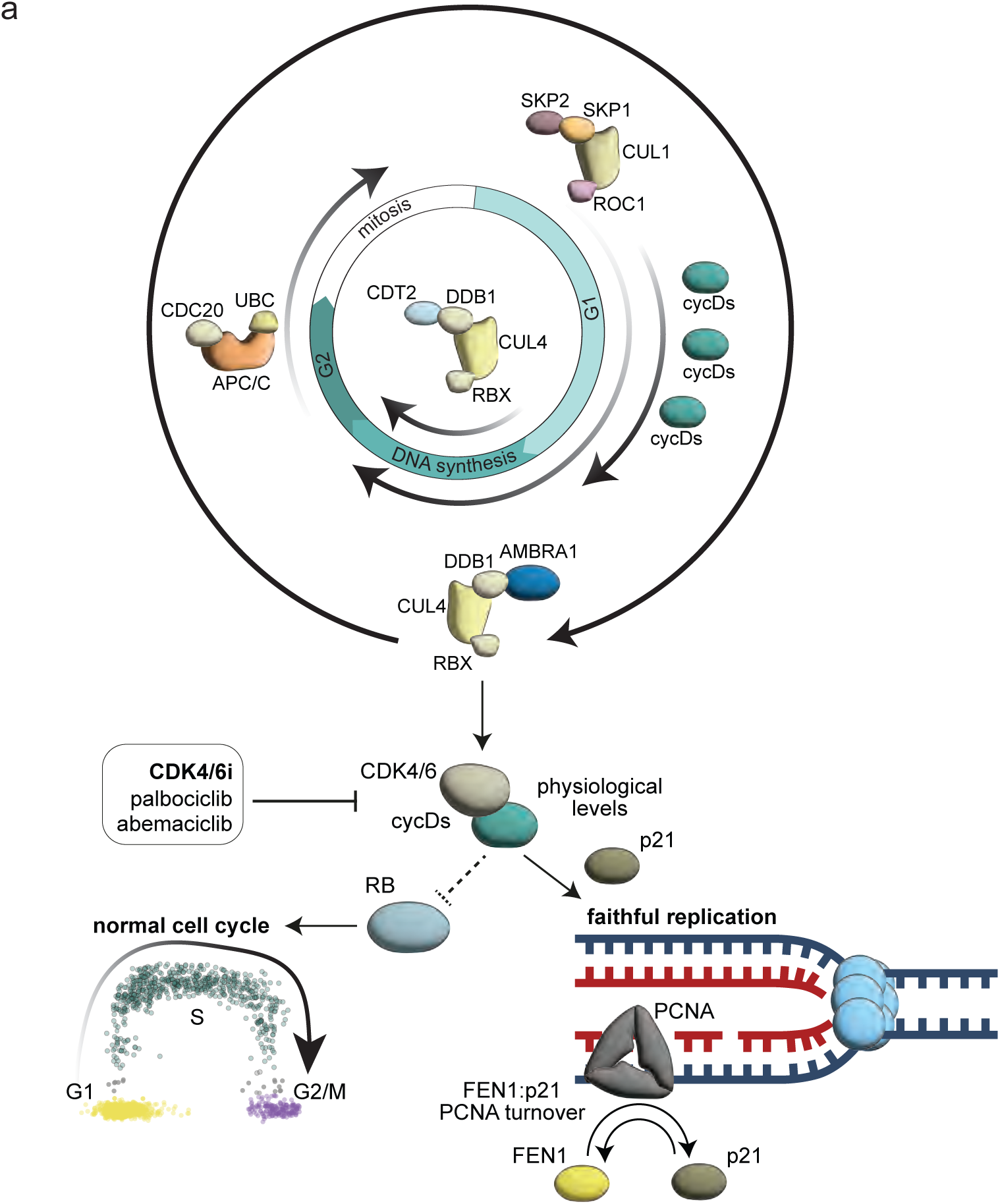
Interplay between AMBRA1-Dependent p21 degradation and other E3 Ligases in cell cycle regulation and PCNA processivity. **a,** Graphical representation illustrating the various pathways involved in the degradation of p21 and D-type cyclins (cycDs). The substrate receptor AMBRA1 plays a key role in targeting these proteins for degradation via the CRL4-DDB1 complex. This regulation occurs throughout the different phases of the cell cycle, ensuring proper checkpoint transition and accurate DNA replication by maintaining physiological levels of the CDK4/6–cyclin D–p21 complex.

